# MIND WANDERING DURING IMPLICIT LEARNING IS ASSOCIATED WITH INCREASED PERIODIC EEG ACTIVITY AND IMPROVED EXTRACTION OF HIDDEN PROBABILISTIC PATTERNS

**DOI:** 10.1101/2024.07.30.605743

**Authors:** Péter Simor, Teodóra Vékony, Bence C. Farkas, Orsolya Szalárdy, Tamás Bogdány, Bianka Brezóczki, Gábor Csifcsák, Dezső Németh

**Author notes:** **Conflict of interest:** The authors have declared that no conflict of interest exists.

## Abstract

Mind wandering, occupying 30-50% of our waking time, remains an enigmatic phenomenon in cognitive neuroscience. Predominantly viewed negatively, mind wandering is often associated with detrimental impacts on attention-demanding (model-based) tasks in both natural settings and laboratory conditions. Mind wandering however, might not be detrimental for all cognitive domains. We proposed that mind wandering may facilitate model-free processes, such as probabilistic learning, which relies on the automatic acquisition of statistical regularities with minimal attentional demands. We administered a well-established implicit probabilistic learning task combined with mind wandering thought probes in healthy adults (N = 37, 30 females). To explore the neural correlates of mind wandering and probabilistic learning, participants were fitted with high-density electroencephalography. Our findings indicate that probabilistic learning was not only immune to periods of mind wandering, but was positively associated with it. Spontaneous, as opposed to deliberate mind wandering, was particularly beneficial for extracting the probabilistic patterns hidden in the visual stream. Additionally, cortical oscillatory activity in the low-frequency (slow and delta) range, indicative of covert sleep-like states, was associated with both mind wandering and improved probabilistic learning, particularly in the early stages of the task. Given the importance of probabilistic implicit learning in predictive processing, our findings provide novel insights into the potential cognitive benefits of task-unrelated thoughts in addition to shedding light on its neural mechanisms. This surprising benefit challenges the predominant view of mind wandering as solely detrimental and highlights its complex role in human cognition, especially in memory consolidation.

**Statement of significance:** Mind wandering poses an unresolved puzzle for cognitive neuroscience: it is associated with poor performance in various cognitive domains, yet humans spend 30-50% of their waking time mind wandering. We proposed that mind wandering may be beneficial for less attention-demanding cognitive processes requiring automatic, habitual learning. We assessed an implicit probabilistic learning task measuring the ability to extract (without awareness) hidden regularities from the information stream. Participants showed superior performance in probabilistic learning during periods of mind wandering, especially when such task-unrelated thoughts occurred spontaneously without intention. Moreover, mind wandering and probabilistic learning were both associated with slow frequency neural activity, suggesting that mind wandering may reflect a transient, offline state facilitating rapid learning and memory consolidation.

## Introduction

Cognitive control is essential for maintaining focus on goal-related information and enabling efficient motor responses, thereby improving performance in a variety of laboratory tasks. Conversely, losing focus and experiencing mind wandering diminishes performance in areas such as sustained attention (Smallwood et al., 2004), executive control (McVay and Kane, 2010, 2012a, 2012b; Andrillon et al., 2021), reading comprehension (Bonifacci et al., 2023), and explicit sequence learning (Brosowsky et al., 2021).

Mind wandering; however, may not be detrimental to behavior in all cases. Broadening the scope of attention and leveraging spontaneous, unconstrained thoughts could be beneficial, especially under cognitively less demanding task conditions, or when the goals and rules of the task are not clear (Thompson-Schill et al., 2009; Amer et al., 2016). One example may be statistical (probabilistic) learning, which allows the unintentional extraction of probabilistic regularities of the environment through mere exposure and unsupervised practice (Fiser and Aslin, 2001; Aslin, 2017; Pedraza et al., 2024). Notably, in probabilistic learning tasks, participants automatically acquire stimulus-outcome dependencies without explicit awareness, cognitive control, or focused attention.

Recent studies suggest that probabilistic learning may not only be unaffected by the disruptive effects of mind wandering but could also derive particular benefits from it. Decker and colleagues (2023) observed that in a modified flanker task, where distracting items had a hidden probabilistic relationship with the target items, lapses of attention (estimated from response times) facilitated the learning of these apparently goal-irrelevant probabilistic contingencies (Decker et al., 2023). Nonetheless, mind wandering was not directly assessed in this study. The association between mind wandering and probabilistic learning was directly examined by an online study using the Alternating Serial Reaction Time (ASRT) task embedded with thought-probes on mind wandering (Vékony et al., 2023). In the ASRT task (Howard Jr and Howard, 1997), implicit statistical learning indicating predictive processes and visuomotor learning are measured simultaneously. Whereas the former reflects the acquisition and anticipation of probabilistic stimulus-response dependencies, the latter captures the efficiency of visuo-spatial discrimination (Vékony et al., 2023). Notably, while participants aim to provide accurate motor responses to a stream of visual inputs, they unintentionally and without awareness extract the probabilistic patterns hidden in the visual stream. This study revealed that participants exhibited improved probabilistic learning but reduced visuomotor accuracy when they reported mind wandering, indicating that mind wandering despite its evident costs may facilitate the extraction of predictable patterns of the environment (Vékony et al., 2023).

The underlying mechanisms of mind wandering; however, still remain unexplored. Theoretical speculations suggest that attenuated sensory processing during mind wandering shares some resemblance with states of sleep (Andrillon et al., 2019; Jubera-Garcia et al., 2021; Simor et al., 2023). Moreover, mind wandering may represent a short-lasting, offline state (i.e. transiently decoupled from the external inputs) that enhances the processing and consolidation of previously encoded material, resembling the mechanisms of sleep-dependent memory consolidation (Jubera-Garcia et al., 2021; Wamsley, 2022; Vékony et al., 2023). Accordingly, mind wandering has been linked to decreased evoked potentials, indicating diminished sensory processing (Kam et al., 2022), as well as increased, slow-frequency activity reminiscent of the slow and delta waves observed during non-rapid eye movement (NREM) sleep (Andrillon et al., 2021; Wienke et al., 2021). In addition, sleep-like cortical activity during mind wandering was expressed in a region-specific, topographically localized manner (Andrillon et al., 2021; Wienke et al., 2021), pointing to the links between mind wandering and local sleep (Andrillon et al., 2019). The electroencephalographic (EEG) correlates of mind wandering, however, were not limited to slow frequency activity, but also to increases in theta and alpha power (Kam et al., 2022).

Here, we aimed to i) provide additional support for the benefit of mind wandering in probabilistic learning, ii) explore the neural correlates of mind wandering during task performance focusing on aperiodic and periodic components of the EEG spectra (Donoghue et al., 2020), and iii) examine if mind wandering and probabilistic learning shares neural features in terms of overlapping frequency- and scalp-specific characteristics.

## Methods

### Participants

Thirty-seven participants (Mean age: 22.1 years ± 1.27, 30 females) were recruited from university students who volunteered for scientific experiments in exchange for partial course credits. Participants had no history of psychiatric, neurological, or chronic somatic disorders, were not currently using medication that could affect alertness, mood, cognition, or sleep, and had normal or corrected-to-normal vision. Only, right-handed participants were included in the study as verified by the Edinburgh handedness inventory (Oldfield, 1971). Participants were not informed about the experiment’s purpose until after completing the tasks; however, they were debriefed afterward. The study received approval from the Research Ethics Committee of Eötvös Loránd University, and all participants provided informed consent.

### Procedure

Participants arrived at the laboratory between 12.00 and 14.00. They were fitted with 64-channel scalp electrodes (see below) and received detailed instructions about the study protocol. They were first introduced to the Alternating Serial Reaction Time (ASRT) task, which involved pressing the key corresponding to the direction of the target stimulus as quickly and accurately as possible, using their left middle and index fingers and their right index and middle fingers. Participants were informed that after each block of the ASRT task, they would be asked three questions to evaluate their thoughts in the previous block and assess their level of mind wandering (MW) which was operationalized as task-unrelated perceptions, thoughts, or memories. A detailed explanation was provided on the different options along with examples of how participants should respond in various scenarios (see **S1** in **Supplementary Materials**). Then, participants completed a short quiz to evaluate their understanding of how to answer the questions about their thoughts, with feedback and explanations provided afterward. Participants had the option to retake the quiz or proceed to the task (see **S2 Supplementary Materials** for details on the quiz). Following the two initial practice blocks of the ASRT task with random stimuli, participants completed 30 additional blocks of the ASRT task. After responding to the thought probes in each block, participants received performance feedback, which included information on both mean speed and accuracy. To guarantee that the ASRT task functioned in a similar manner to previous studies, we assessed the participants’ conscious knowledge of the hidden sequence following previously validated protocols (Horváth et al., 2022; Vékony et al., 2022). Participants were asked if they noticed anything unusual or any regularities in the task, and if so, to elaborate on their response. None of the participants were able to accurately describe the alternating sequence. After completing the ASRT task, participants were asked to respond to demographic questions (age, gender, education, etc.). One participant had missing data due to technical issues in 10 blocks (16-25 blocks), but their remaining data was retained in the analyses.

### The ASRT task

The ASRT task embedded with thought probes was displayed on a 24-inch monitor screen in a sound-attenuated room. Responses to the task were recorded using a Cedrus RB-530 response pad (Cedrus Corporation, San Pedro, CA). A modified version of the task suitable for EEG analyses (Kóbor et al., 2018) was used to measure Probabilistic Learning and Visuomotor Performance. In this task version, an arrow stimulus appeared at the center of the screen, and participants were instructed to press the corresponding key to the spatial direction (up, down, left, or right) of the arrow as accurately and as fast as they could. The arrow was presented for 200 ms, followed by the presentation of a fixation cross for 500 ms during which the participant could respond. Subsequently, a fixed delay of 750 ms followed, displaying again a fixation cross, and then, the next trial appeared (applying fixed inter stimulus intervals). In case of a wrong response, a “×” appeared in the middle of the screen for 500 ms, and in case of a missing response, an “!” appeared for the same duration. Following this, a fixation cross appeared for 250 ms before the next trial began. The stimuli followed a probabilistic eight-element sequence, alternating between patterned and random elements (e.g., 2 – R – 4 – R – 3 – R – 1 – R, where ‘R’ indicates a random direction and the numbers represent predetermined directions from up, down, left, and right). Each participant was assigned one of 24 possible sequences, which they encountered throughout the task. The ASRT task comprised 30 blocks, each consisting of ten repetitions of the eight-element sequence (80 trials) preceded by 5 random trials at the beginning of each block for warm-up purposes. Following each block, participants took a brief break and were instructed to respond to the mind wandering thought probes before resuming the task (**Fig 1B**). The ASRT task incorporated a probabilistic sequence structure where certain runs of three consecutive stimuli (triplets) appeared with a higher probability (high-probability triplets) than others (low-probability triplets). A trial refers to a single element in the sequence, which could be either a pattern or random element and, importantly, also the last element in a high-or low-probability triplet. It is crucial to note that the analysis focuses on whether the provided trial constitutes the final element of a high-or low-probability triplet, rather than its classification as a pattern or random element within the alternating sequence. For instance, in a sequence like 2 – R – 4 – R – 3 – R – 1 – R, triplets such as 2-X-4, 4-X-3, 3-X-1, and 1-X-2 (where X represents the middle element of a triplet) occurred more frequently than triplets like 2-X-1 or 2-X-3 (see **Fig. 1C**). It is important to highlight that when referring to triplet type later on, the focus is on trials serving as the final element of a high- or low-probability triplet. Throughout the task, a total of 64 distinct triplets could potentially occur (16 with high probability and 48 with low probability). High-probability triplets could be formed either by having two pattern trials and one random trial in the center (occurring in 50% of trials) or by having two random trials and one pattern trial in the center (occurring in 12.5% of trials). Among all trials, 62.5% represented the last element of a high-probability triplet, while 37.5% were assigned to the last element of a low-probability triplet (see **Fig. 1D**).

**Figure 1.**
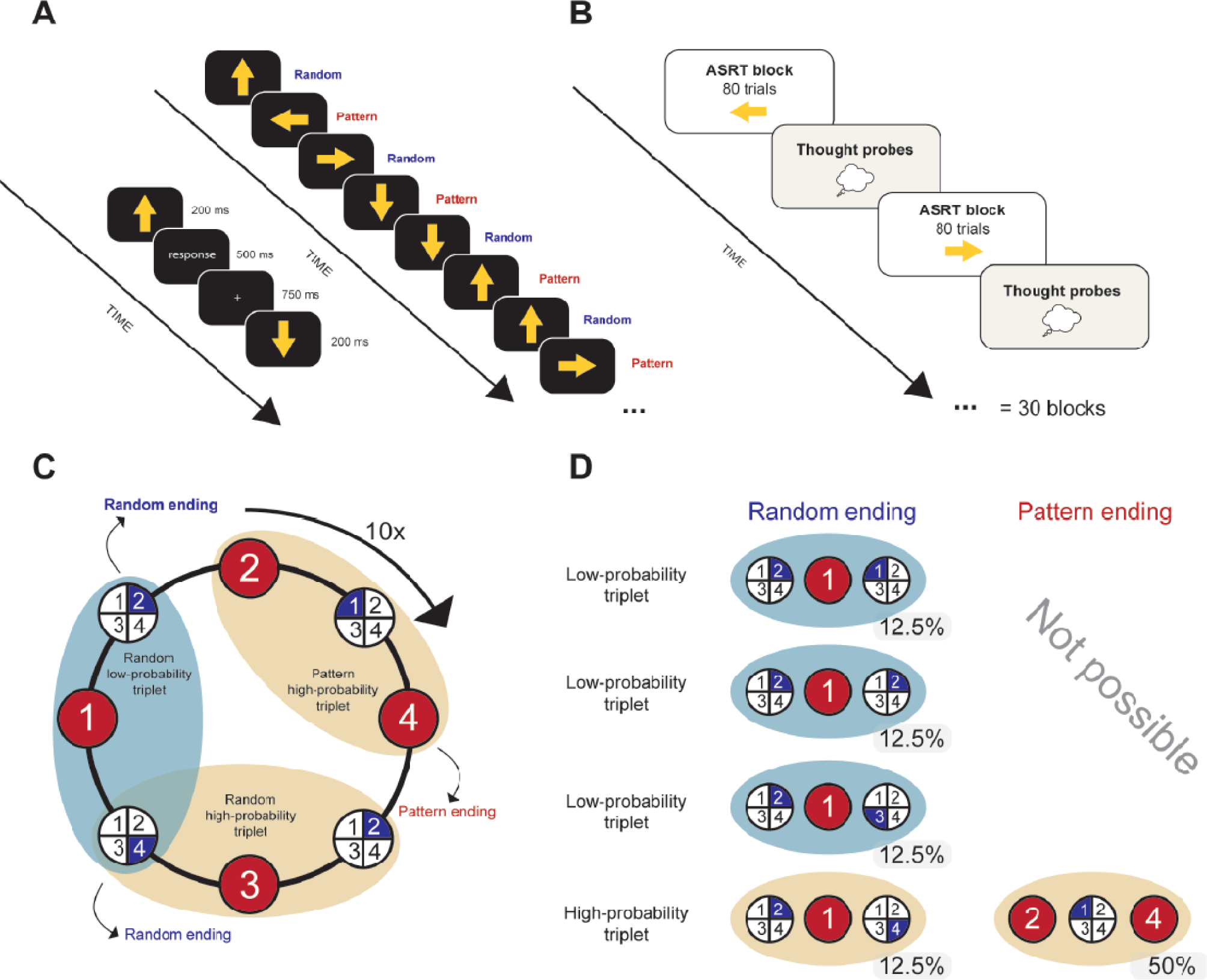
Experimental design and task structure of the ASRT task. **(A)** In the ASRT task, participants had to press keys corresponding to the direction of an arrow that appeared in the center of the screen. Every second trial was part of an 8-element probabilistic sequence. Random elements were inserted among pattern elements to form the sequence (e.g., 2-R-4-R-3-1-R, where numbers indicate the direction of pattern trials (up, down, left, or right), and *r* represents random directions of the four possible directions). **(B)** The experiment consisted of 30 blocks with mind wandering thought probes administered after each block of 80 trials. Participants were asked to reflect on their thoughts during the block and decide whether the last block was dominated by 1) mind wandering versus task focus, 2) mind wandering versus mind blanking, and 3) spontaneous versus deliberate mind wandering. **(C)** Formation of triplets in the task. Pattern elements are represented by red backgrounds (they are constantly pointing at that direction throughout the task), and random elements are represented by blue backgrounds (they are always chosen from the four possible directions randomly). Every trial was categorized as the third element of three consecutive trials (a triplet). It is worth highlighting that the analysis depends on whether the given trial is the last element of a high-or low-probability triplet, rather than its classification as a pattern or random element within the alternating sequence. The probabilistic sequence structure resulted in a higher occurrence of some triplets (high-probability triplets) than others (low-probability triplets). Please note that only three triplets are highlighted on this subfigure for visualization purposes [2(P)-1(R)-4(P) as a pattern-ending high-probability triplet, 2(R)-3(P)-4(R) as a random-ending high probability triplet, and 4(R)-1(P)-2(R) as a random-ending low-probability triplet]. However, every three consecutive elements form either a high-or low-probability triplet. Therefore, in the above example, from these 8 consecutive elements - 2(P)-1(R)-4(P)-2(R)-3(P)-4(R)-1(P)-2(R) - 6 triplets can be formed. If we consider the pattern element 2 as a starting point, then the triplets are in the following order: 2(P)-1(R)-4(P), 1(R)-4(P)-2(R), 4(P)-2(R)-3(P), 2(R)-3(P)-4(R), 3(P)-4(R)-1(P) (these are all high-probability triplets), and 4(R)-1(P)-2(R) (low-probability triplet). **(D)** The formation of high-probability triplets could have involved the occurrence of either two pattern trials and one random trial at the center, which transpired in 50% of trials, or two random trials and one pattern trial at the center, which occurred in 12.5% of trials. In total, 62.5% of all trials constituted the final element of a high-probability triplet, while the remaining 37.5% were the final elements of a low-probability triplet.

### Mind wandering thought probes

Following each block of the ASRT task, participants were prompted to reflect on their thoughts during the last block and respond to questions regarding their attentional states (**Fig.1B**). More specifically, they were asked to rate if in the last block 1) they engaged in mind wandering or maintained task focus; 2) in case they experienced mind wandering they were thinking of something particular, or did not think about anything (“mind blanking”); and 3) whether their attentional state was controlled deliberately or was rather spontaneous. This way, after each block, participants were presented with three items to be rated on a four-point scale: Q1) *To what degree were you focusing on the task just before this question?* (1 - Not at all; 4 - Completely), Q2) *To the degree to which you were not focusing on the task, were you thinking of something in particular or just thinking about nothing?* (1 - I was thinking about nothing, 4 - I was thinking about something in particular), and Q3) *Were you deliberate about where you focused your attention (either on-task or elsewhere) or did it happen spontaneously?* (1 - I was completely spontaneous, 4 - I was completely deliberate). While the first question has been used in prior studies (Alexandersen et al., 2022; Groot et al., 2022), the latter two were tailored to differentiate between MW with reportable content versus MB (Q2), and between spontaneous versus deliberate MW (Q3). Although it is common to directly distinguish between on-task periods and either MW versus MB, or unintentional versus intentional MW with specific questionnaires (Seli et al., 2018), we chose to explore these aspects of MW in two follow-up questions to avoid overwhelming participants with numerous response options and to gain a more nuanced understanding of their mental states during the ASRT task (See Aasen et al., 2024; Drevland et al., 2024).

### EEG recording and preprocessing

EEG activity was measured by a 64-channel recording system (BrainAmp amplifier and BrainVision Recorder software, BrainProducts GmbH, Gilching, Germany). The Ag/AgCl sintered ring electrodes were mounted in an electrode cap (EasyCap GmbH, Herrsching, Germany) on the scalp according to the 10% equidistant system. During acquisition, electrodes were referenced FCz electrode but were re-referenced to the average of the mastoids before further data processing. Horizontal and vertical eye movements were monitored by EOG channels. EMG electrodes to record muscle activity were placed on the chin. All electrode contact impedances were kept below 10 kΩ. EEG data was recorded with a sampling rate of 1000 Hz. Antialiasing digital filters and notch filters to remove power line noise were applied. Preprocessing of EEG data was performed by using custom-made scripts in MATLAB (version 9.14.0.2206163, R2023a, The Mathworks INC, Natick, MA) as well as functions of the Fieldtrip toolbox (Oostenveld et al., 2011). EEG recordings were band-pass filtered between 0.3 and 70 Hz (with Butterworth, zero phase forward and reverse digital filters). We performed independent component analysis (ICA) to identify cardiac, eye movement, and other muscular artifacts using FieldTrip routines (Oostenveld et al., 2011). Independent components, primarily two to three (up to four maximum), representing eye movements and muscular artifacts, were semi-automatically detected and identified by inspecting waveforms and their topographical distribution (Campos Viola et al., 2009). Additionally, a semi-automatic artifact rejection tool was applied using the ft_artifact_zvalue function of the FieldTrip routine. This procedure involves filtering the data and averaging it across channels after Z-transformation. Accumulated z-scores are then visualized, and thresholds are flexibly set and adapted to each recording. Subsequently, trials identified as artifacts were visually inspected to confirm and discard those with technical or movement-related artifacts.

### Data analyses I: Behavioral and thought probes

We used MATLAB to process behavioral data assessed in the ASRT task. Each trial (representing an arrow pointing to one of the four directions) was categorized based on the two preceding trials as the last element of a high-or low-probability triplet. Trills (e.g., 1-2-1), repetitions (e.g., 2-2-2), the first two trials, and trials with a reaction time above 1000 ms were removed from the analysis. Inaccurate responses were also removed from the analyses of RTs. We defined two types of behavioral measures: *Probabilistic Learning* and *Visuomotor Performance*. *Probabilistic Learning* was operationalized as the difference in RT/accuracy between high-probability and low-probability trials (i.e., between the third element of a high-probability triplet and the third element of a low-probability triplet. *Visuomotor Performance*, on the other hand, was operationalized as the overall RT/accuracy on the task and its changes over time (i.e., lower accuracy in later blocks) regardless of the trial probability. *Probabilistic Learning* and *Visuomotor Performance* scores were extracted considering the mean accuracy scores (percentage of correct responses/all trials) and the median of RTs of accurately responded trials and for each block. Similarly, the same measures were extracted for each time bin comprising the averages for chunks of 5 consecutive blocks (Block 1-5, 6-11…25-30). Since mind wandering was previously linked to accuracy based measures of probabilistic learning (Vékony et al., 2023), we focused on this metric, but also examined RT based measures (reported in the **Supplementary Materials**.) Responses to mind wandering thought probes were similarly extracted for each block and time bin. For time bins, responses to thought probes were averaged for each chunk of 5 consecutive blocks. Scores reflecting *Mind Wandering* versus *Task Focus* were present for each block. However, scores representing *Mind Wandering* versus *Mind Blanking*, and *Spontaneous* versus *Deliberate* mind wandering were only considered when participants reported a tendency to mind wander (responses 1 and 2) instead of focusing on the task (responses 3 and 4) during the block (i.e., to Q1).

### Data analyses II: EEG analyses

EEG analyses were performed by using custom-made scripts and Fieldtrip routines in MATLAB. We extracted the aperiodic and periodic spectral components of EEG activity measured during the 30 blocks of the ASRT task. To quantify the spectral slope as well as the oscillatory (periodic) activity of the EEG signal we applied the FOOOF (fitting oscillations and one over f) method (Donoghue et al., 2020). Separating the EEG spectra into aperiodic and periodic components is an efficient technique to delineate if changes in power are caused by a shift in aperiodic background activity following a power law or increases/decreases in oscillatory activity beyond the aperiodic background. Moreover, the parametrization of aperiodic and periodic activity reduces the redundancy of the power spectrum featuring largely correlated power values across different frequencies (Donoghue et al., 2020; Bódizs et al., 2021; Gerster et al., 2022; Schneider et al., 2022). Parametrization of the EEG spectra proved to be a sensitive approach to track different states of consciousness (Colombo et al., 2019; Bódizs et al., 2021), cognitive performance (Ouyang et al., 2020), and shifts in modality-specific attention (Waschke et al., 2021).

Artifact-free EEG data segmented into 2-s long non-overlapping time windows was subjected to Fast Fourier Transformation (FFT) using discrete prolate spheroidal sequences (DPSS) multitapers to attenuate spectral leakage. Power spectral densities (PSD) between 1.5 and 40 Hz with 0.5 Hz resolution were obtained, and were averaged over the segments within each block and each participant. Hence, the power spectra were used as the input to extract the aperiodic and periodic components. In brief, the FOOOF method computes a linear fit to the spectrum represented in a log-log scale. This linear fit is then subtracted from the spectrum yielding a flattened spectrum. Using the flattened spectrum a relative and an absolute threshold is set to two times the standard deviation of the flattened spectrum. Next, the method fits Gaussian functions to the largest peak exceeding the threshold, subtracts it from the spectrum, and then iteratively proceeds to the next largest peak until no more peaks surpassing the threshold are detected. The oscillatory components are then derived by fitting a multivariate Gaussian to all extracted peaks simultaneously. Following these iterations, the initial fit is reintegrated into the peak-free flattened PSD, resulting in the aperiodic component of the PSD. Subsequently, this aperiodic component undergoes another fitting process, producing the final fit with parameters for y-intercept and slope. We used the spectral slope in our analyses, which is assumed to reflect neural inhibitory and excitatory balance (Gao et al., 2017; Donoghue et al., 2020), with larger values (steeper slopes) reflecting stronger inhibition. Periodic activity was inferred from the flattened PSD. Band-wise oscillatory activity was obtained by averaging power values between 1.5-2 Hz (Slow), 2.5-4 Hz (Delta), 4.5-8.5 Hz (Theta), 9-13.5 Hz (Alpha), and 14-30Hz (Beta), but bin-wise values were also retained for further analyses.

### Statistical Analyses

Statistical analyses and data visualizations were performed in Rstudio (Version 1.4 1717; Allaire, 2012) and MATLAB. Since multiple block-wise values for behavioral, self-report, and EEG variables were nested within each individual, Linear Mixed Models (LMMs) were employed to examine within-person associations between our variables of interest allowing for random effects (random intercept and slope) by participants. For each LMM, we first fitted the maximal random-effects structure (i.e., including random effects for all variables and allowing correlations between them), and then reduced it to achieve convergence by removing random slopes and maintaining random intercepts only. To focus on within-person associations, the numerical predictors were mean-centered (to the individual’s mean values) (Wang and Maxwell, 2015). The relevant subsections of the Results section provide detailed descriptions of the specific outcome variables, as well as the fixed and random effects to ease the understanding of the specific models. In brief, separate LMMs were used to i) examine the rate of *Probabilistic Learning* and *Visuomotor Performance* as a function of practice throughout the successive blocks; ii) to examine within-person fluctuations in self-reported mind wandering throughout the successive blocks; iii) to examine within-person associations between self-reports of mind wandering and behavioral performance (*Probabilistic Learning* and *Visuomotor Performance*) in the ASRT task; iv) and to examine the association of block-wise within-person fluctuations in aperiodic and periodic EEG activity with mind wandering and probabilistic learning. Separate LMMs were run to test the associations of EEG measures (outcome variables) with mind wandering and probabilistic learning (predictors in separate models). In order to control the confounding effect of time on EEG activity, *Block* was entered as a continuous, mean-centered predictor in addition to probe responses about attentional focus (mind wandering versus task focus), as well as in addition to probabilistic learning in the LMMs with EEG measures as outcome variables. To explore the region-specific, topographical aspects of these associations, LMMs were performed relating mind wandering and probabilistic learning with EEG measures at each electrode location. To address the issue of multiple comparisons p-values were adjusted with FDR (false discovery rate) correction (Benjamini and Hochberg, 1995). An alpha value of 0.05 was used for all statistical analyses.

## Results

### The tendency to mind wander fluctuates and gradually increases throughout the task

First, we examined whether the tendency of mind wandering changed throughout the task. Inter- and intra-individual variability and the relative frequencies of *Mind Wandering* vs. *Task Focus* self-reports are depicted in **Fig 2**. Mind wandering was highly variable across the successive blocks showing considerable inter-and intra-individual variability (ICC = 0.45) (**Fig 2A**). Participants tended to mind wander more at later stages of the task (**Fig 2B & C**). This was revealed by a LMM (with random intercepts and slopes by participant) regressing self-reports of *Mind Wandering* vs. *Task Focus* (outcome variable on a 4-point scale for thought probe Q1) on *Block* number (predictor), as indicated by the main effect of *Block* (b = −0.13, 95 % CI = [-0.19, −0.06], t = −4.19, p < 0.001). Likewise, a GLMM regressing dichotomized *Mind Wandering* vs. *Task Focus* scores (binary outcome variable) showed a similar main effect of *Block* (Odds Ratio = 0.69, 95% CI = [0.61, 0.78], t = −6.3, p < 0.001). To study if the phenomenological aspects of MW experiences varied throughout the task, we also examined if *Mind Blanking vs. Mind Wandering* and *Spontaneous vs. Deliberate MW* were predicted by *Block. Block* number as a predictor in LMM (with random intercepts and slopes by participant) did not significantly predict the tendency of *Mind Blanking vs. Mind Wandering* (b = 0.04, 95% CI = [-0.13, 0.22], t = 0.47, p = 0.64) or *Spontaneous vs. Deliberate MW* (b = −0.03, 95% CI = [-0.17, 0.11], t = −0.44, p = 0.66).

**Figure 2.**
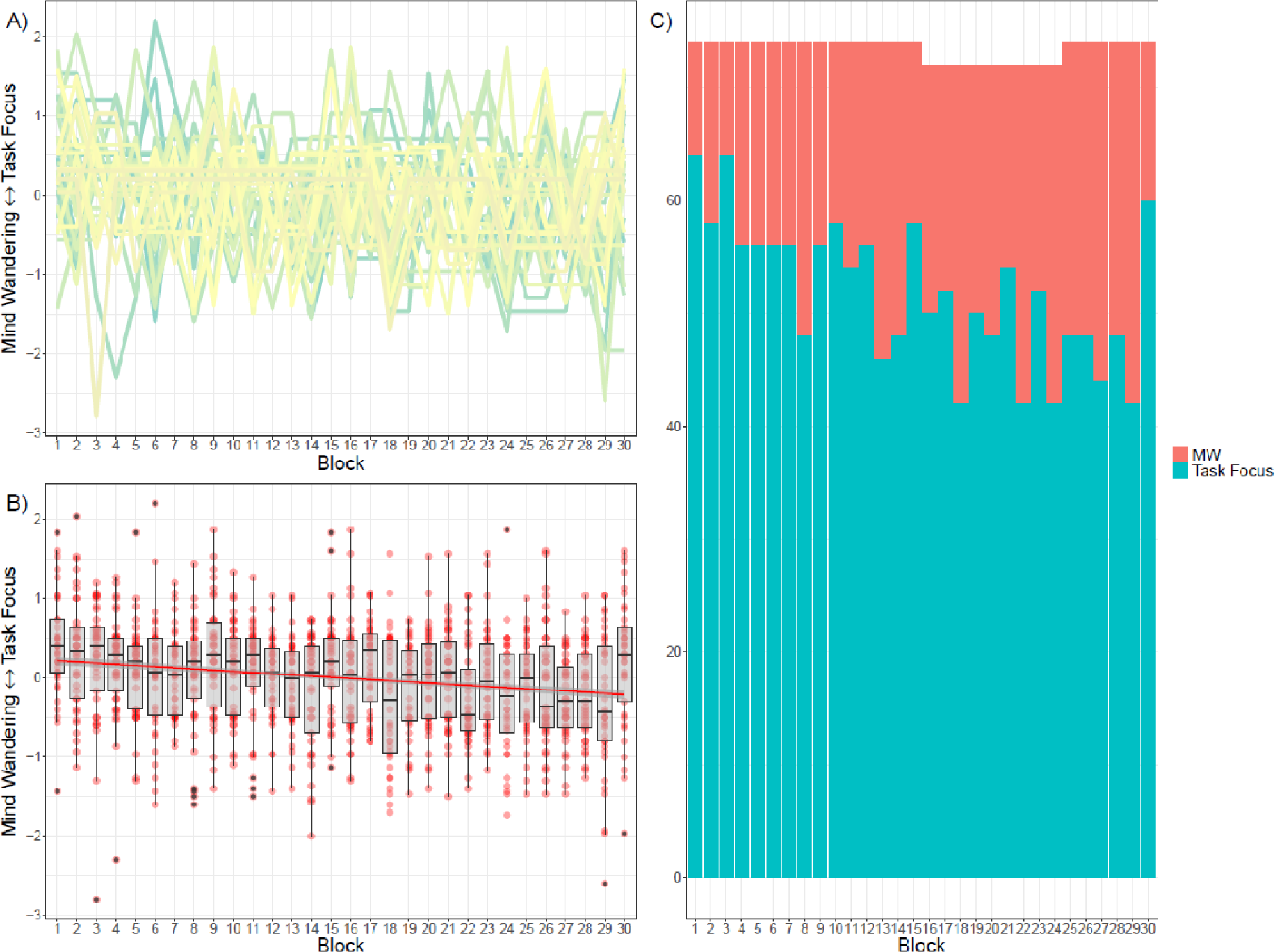
Variability and temporal trajectory of mind wandering during the ASRT task. A) Mind wandering exhibits considerable variability across and within individuals throughout the successive blocks. Each line (distinguished by different colors) represents different individual’s responses over the blocks. Values on the y-axis are centered for visualization only. B) *Mind Wandering* vs. *Task Focus* scores over the successive blocks. Self-reports of mind wandering are gradually more prevalent as the task progresses. Values were centered block-wise to the overall individual means for visualization purposes. Lower scores on the Y-axis indicate increased mind wandering. C) The amount of *Mind Wandering* vs. *Task Focus* states by dichotomized values. Besides the final block, mind wandering becomes more prevalent over the course of the task.

### Mind wandering is linked to better probabilistic learning but poorer visuomotor accuracy

To investigate if mind wandering was associated with acquiring hidden probabilistic regularities and with overall visuomotor performance, we entered the predictors *Block*, *Mind Wandering vs. Task Focus scores*, as well as their interaction, on the outcome measures of (accuracy and RT based) *Probabilistic Learning* and *Visuomotor Performance (*termed as *Visuomotor Accuracy* and *Visuomotor RT),* in separate LMMs. With regards to accuracy, *Probabilistic Learning* was defined as the difference in block-wise responses to high-versus low-probability trials, whereas *Visuomotor Accuracy* represented overall, block-wise accuracy regardless of trial types. First, we evaluated if learning occurred over the course of the task. As shown in **Table 1**, a main effect of *Block* emerged indicating increased *Probabilistic Learning* over the course of the task, that is, participants were gradually better in extracting the statistical pattern hidden in the task, becoming more accurate (**Fig 3A&C**) in responding to high-probability versus low-probability trials. *Block* was also significantly, and negatively associated with *Visuomotor Accuracy*, pointing to overall less accurate (**Fig 3B&D)** responses as the task progressed, regardless of the probabilistic nature of trials. Reaction time based measures exhibited a similar learning pattern (see **S3** in **Supplementary Materials**). Participants were faster in responding to high-versus low-probability trials, and showed generally faster reaction times (regardless of trial types) as the task progressed (See **S4** in **Supplementary Materials**.)

**Figure 3.**
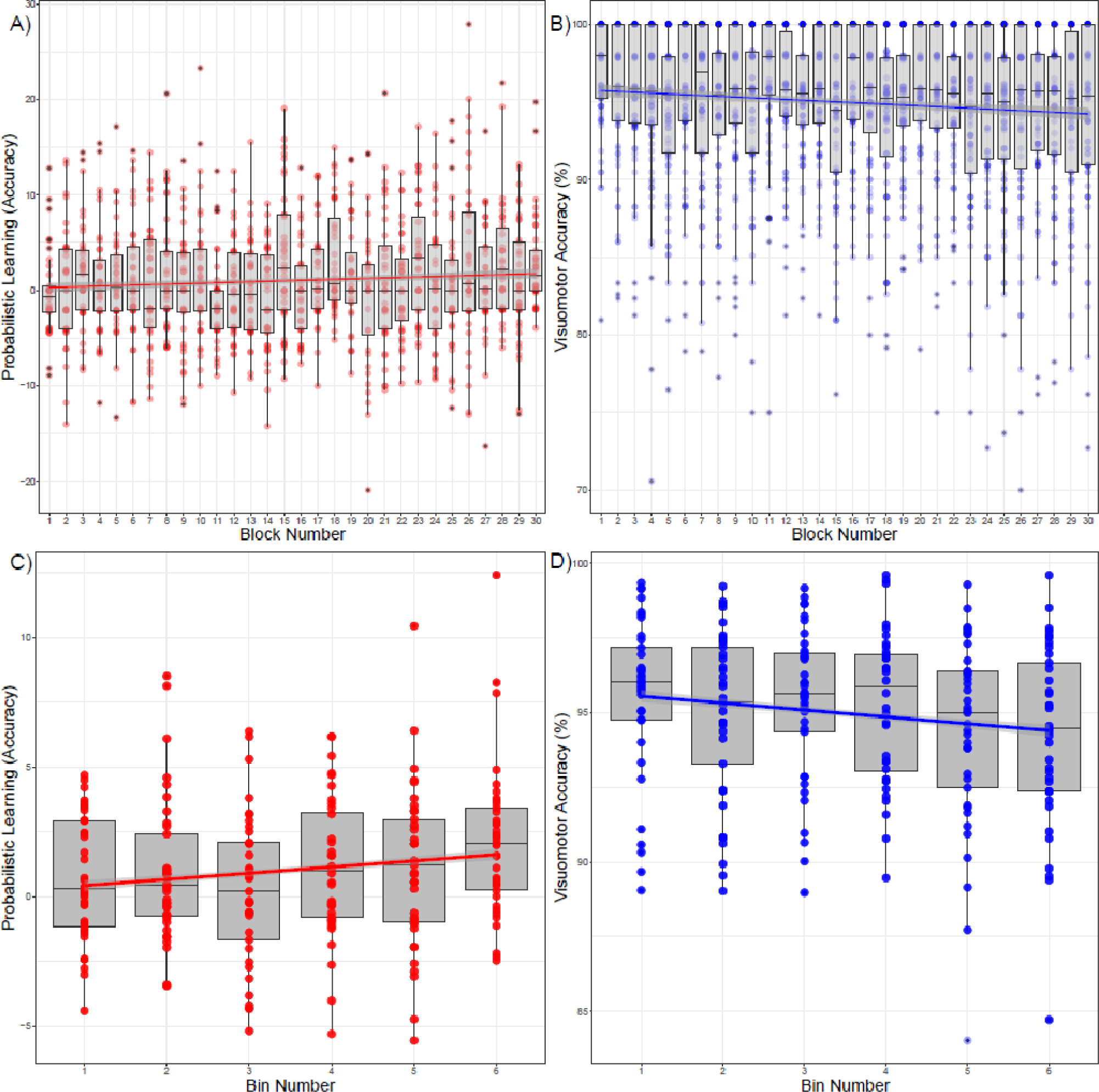
Probabilistic Learning and Visuomotor Accuracy over successive blocks and bins. *Probabilistic Learning* is quantified by the difference in accuracy (A, C) (percentage of correct responses) between high-probability and low-probability trials averaged within each block. *Probabilistic Learning* gradually increases throughout the task due to relatively more accurate responses to high-versus low-probability trials. On the other hand, *Visuomotor Accuracy* (overall accuracy regardless of trial types) (B, D) is gradually reduced as the task progresses, reflecting less accurate responses regardless of trial types. The upper graphs depict performance across each block. In the lower graphs (to facilitate visualization) the same performance is averaged over successive steps of five consecutive blocks (Blocks: 1-5, 6-10…26-30) marked as bins.

**Table 1.**
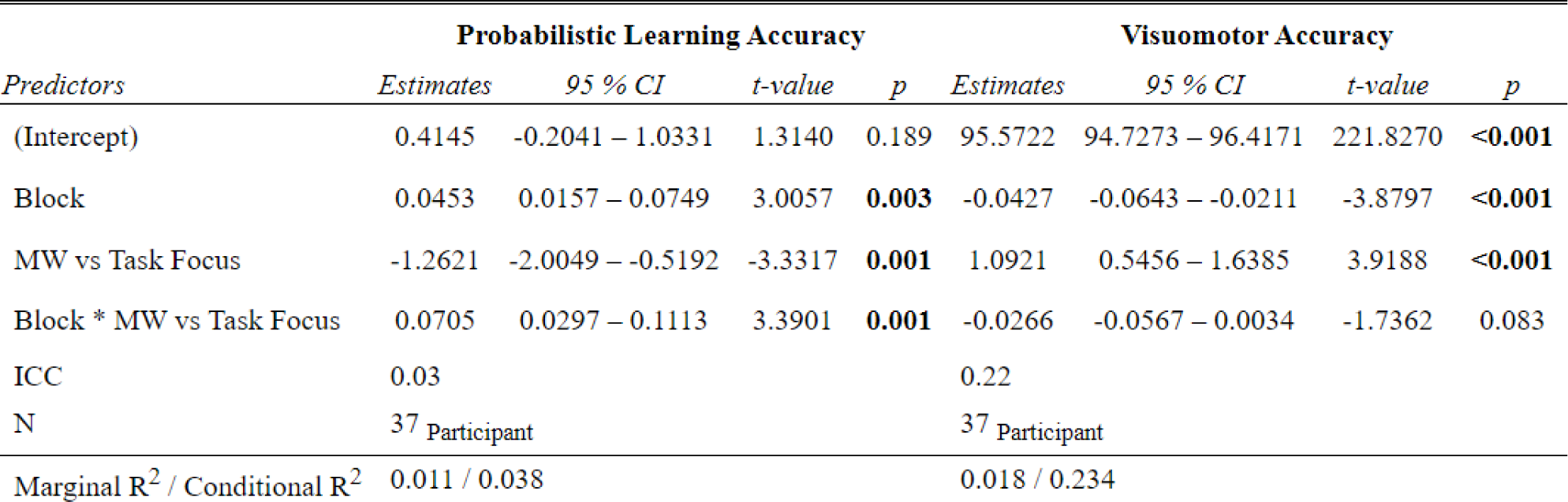
Mind wandering and ASRT task performance. LMM random intercept models with accuracy based indices of *Probabilistic Learning* and *Visuomotor Accuracy* as outcome variables, *Block* and *Mind Wandering* vs. *Task Focus*, as well as their interaction as fixed effects predictors.

| *Predictors* | **Probabilistic Learning Accuracy** |  |  |  | **Visuomotor Accuracy** |  |  |  |
| --- | --- | --- | --- | --- | --- | --- | --- | --- |
|  | *Estimates* | *95 % CI* | *t-value* | *p* | *Estimates* | *95 % CI* | *t-value* | *p* |
| (Intercept) | 0.4145 | -0.2041 – 1.0331 | 1.3140 | 0.189 | 95.5722 | 94.7273 – 96.4171 | 221.8270 | <0.001 |
| Block | 0.0453 | 0.0157 – 0.0749 | 3.0057 | **0.003** | -0.0427 | -0.0643 – -0.0211 | -3.8797 | <0.001 |
| MW vs Task Focus | -1.2621 | -2.0049 – -0.5192 | -3.3317 | **0.001** | 1.0921 | 0.5456 – 1.6385 | 3.9188 | <0.001 |
| Block * MW vs Task Focus | 0.0705 | 0.0297 – 0.1113 | 3.3901 | **0.001** | -0.0266 | -0.0567 – 0.0034 | -1.7362 | 0.083 |
| ICC | 0.03 |  |  |  | 0.22 |  |  |  |
| N | 37 Participant |  |  |  | 37 Participant |  |  |  |
| Marginal R<sup>2</sup> / Conditional R<sup>2</sup> | 0.011 / 0.038 |  |  |  | 0.018 / 0.234 |  |  |  |

Block spans between 1 and 30. In the case of MW vs. Task Focus lower scores indicate relatively increased mind wandering as centered to the participant’s own mean. CI refers to confidence intervals of 95%.

As shown in **Table 1** accuracy based indices of *Probabilistic Learning* were higher during periods when participants reported a tendency to mind wander versus focusing on the task (b = - 1.26, 95 % CI = [−2.00, −0.52], t = - 3.33, p = 0.001), see **Fig 4A.** Moreover, the interaction between *Mind wandering* vs. *Task Focus* with *Block* (b = 0.07, 95 % CI = [0.03 0.11], t = 3.39, p = 0.001) suggested that mind wandering was associated with better probabilistic learning at earlier phases of the task. To explore this interaction in more detail examining if mind wandering exhibits a distinct association with probabilistic learning in the first (blocks 1-15) versus the second (blocks: 16-30) half of the ASRT task, we employed a further LMM with *Mind wandering vs. Task Focus* and *ASRT time* (*first half* (blocks 1-15) vs. *second half* (blocks 16-30) as predictors of *Probabilistic Learning.* We observed a similar interaction between *Mind wandering* and *ASRT time* (first vs. second part of the ASRT task) (b = 0.88, 95 % CI = [0.13 1.63], t = 2.31, p < 0.02), in addition to the main effects of *ASRT time* (b = 0.77, 95 % CI = [0.25 1.28], t = 2.94, p < 0.003) and *Mind wandering vs. Task Focus* (b = −0.62, 95 % CI = [−1.16 −0.09], t = −2.3, p < 0.02) pointing to increased *Probabilistic Learning* when participants experienced more mind wandering, specifically in the first part of the ASRT task. Post-hoc contrasts showed that the associations between mind wandering and probabilistic learning between the first (EMM: 0.71 95% CI [0.21 1.20]) and the second part of the ASRT task (EMM: 1.42 95% CI [0.97 1.97]) were significantly different (Estimate: −0.8, t-ratio = −2.9 p <0.003).

**Figure 4.**
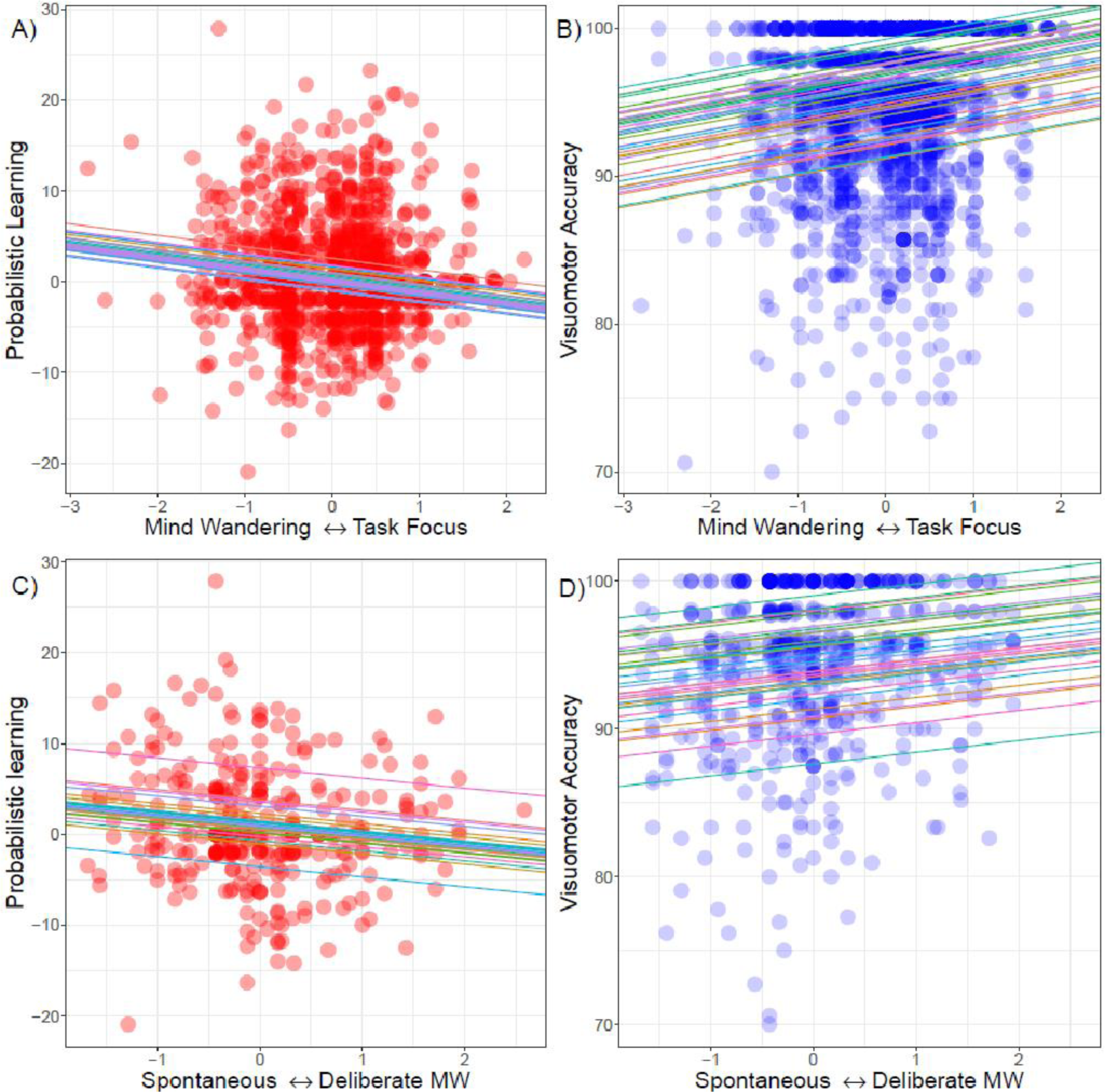
Mind wandering is associated with enhanced Probabilistic Learning and attenuated Visuomotor Accuracy. The results of random intercept and fixed slope LMMs are visualized regressing block-wise centered *Mind Wandering* vs. *Task Focus* scores on (accuracy-based) *Probabilistic Learning* (A) and *Visuomotor Accuracy* (B). Lower values on the X-axes (A & B) indicate increased mind wandering. Similarly, the results of random intercept and fixed slope LMMs are visualized regressing block-wise centered *Spontaneous* vs. *Deliberate Mind Wandering* on (accuracy-based) *Probabilistic Learning* (C) and *Visuomotor Accuracy* (D). Lower values on the X-axes (C & D) indicate increased spontaneous mind wandering. Colored lines represent random intercepts and fixed regression slopes for each participant. Points depict values for the variables of interest of all participants at each block.

With regards to *Visuomotor Accuracy*, *Mind Wandering* vs. *Task Focus* was positively associated with overall accuracy (b = 1.09, 95 % CI = [0.54 1.63], t = 3.91, p < 0.001), indicating that participants committed generally less errors when they tended to focus more on the task (**Fig 4B**). The interaction between *Block* and *Mind Wandering* vs. *Task Focus* was not significant (b = −0.03, 95 % CI = [-0.05 0.00], t = −1.7, p = 0.08). Reaction time based measures of *Probabilistic Learning* and of *Visuomotor Accuracy* were not predicted by self-reports of *Mind Wandering*, or the interaction between *Mind Wandering* and *Block* (See **Table S3** in **Supplementary Materials**).

### The nature of mind wandering relates to probabilistic learning and visuomotor accuracy

We examined if the type of mind wandering was associated with behavioral performance. Therefore, in our next analyses, we considered only those blocks when participants reported to experience mind wandering during the task. *Mind Blanking vs. Mind Wandering* as well as self-reports of *Spontaneous vs. Deliberate mind wandering (MW)* were used as mean-centered predictors of our behavioral measures. Since mind wandering was associated with accuracy based measures of learning, we only considered accuracy based *Probabilistic Learning* and *Visuomotor Accuracy* as our outcome variables. *Mind Blanking vs. Mind Wandering* × *Block* as well as by *Spontaneous vs. Deliberate MW* × *Block* were predictors in separate LMMs (fixed slopes and random intercepts were used for each participant). The experience of *Mind Blanking vs. Mind Wandering* was not associated with *Probabilistic Learning* but mind wandering reports in contrast to mind blanking were associated with better *Visuomotor Accuracy* (See **Table 2** for more details). Notably, more spontaneous (versus deliberate) mind wandering was associated with better *Probabilistic Learning* (**Fig 4C**) and poorer *Visuomotor Accuracy* (**Fig 4D**) (See **Table 2** for more details). The interactions between block number and the nature of mind wandering were not significant on either of the outcome behavioral measures (See **Table 2**).

**Table 2.**
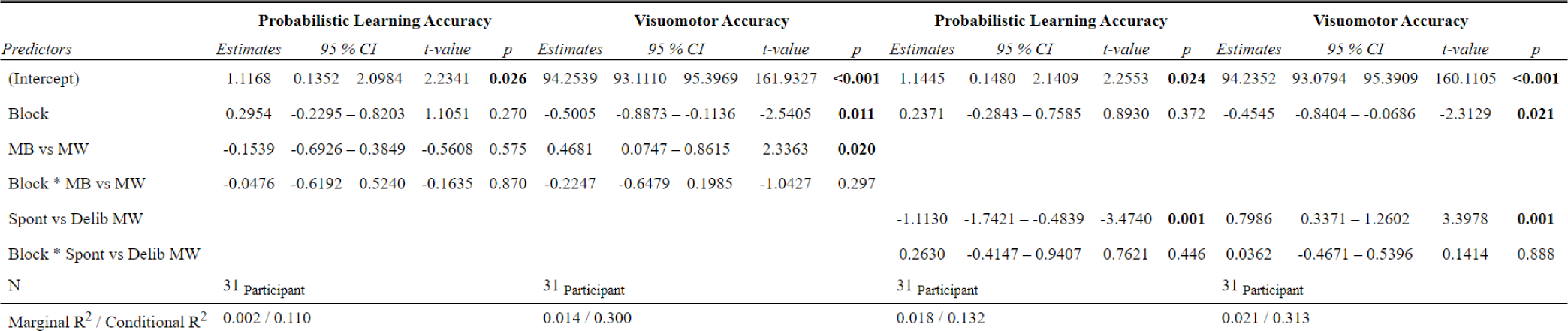
Probabilistic Learning and Visuomotor Accuracy predicted by the subtypes of mind wandering. LMM random intercept models with Probabilistic Learning and Visuomotor Accuracy as outcome variables. Mind Blanking vs. Mind Wandering and Spontaneous vs. Deliberate mind wandering were used as fixed predictors in separate models, taking into account the influence of Block number.

| Predictors | Probabilistic Learning Accuracy |  |  |  | Visuomotor Accuracy |  |  |  | Probabilistic Learning Accuracy |  |  |  | Visuomotor Accuracy |  |  |  |
| --- | --- | --- | --- | --- | --- | --- | --- | --- | --- | --- | --- | --- | --- | --- | --- | --- |
|  | Estimates | 95 % CI | t-value | p | Estimates | 95 % CI | t-value | p | Estimates | 95 % CI | t-value | p | Estimates | 95 % CI | t-value | p |
| (Intercept) | 1.1168 | 0.1352 – 2.0984 | 2.2341 | **0.026** | 94.2539 | 93.1110 – 95.3969 | 161.9327 | **<0.001** | 1.1445 | 0.1480 – 2.1409 | 2.2553 | **0.024** | 94.2352 | 93.0794 – 95.3909 | 160.1105 | **<0.001** |
| Block | 0.2954 | -0.2295 – 0.8203 | 1.1051 | 0.270 | -0.5005 | -0.8873 – -0.1136 | -2.5405 | **0.011** | 0.2371 | -0.2843 – 0.7585 | 0.8930 | 0.372 | -0.4545 | -0.8404 – -0.0686 | -2.3129 | **0.021** |
| MB vs MW | -0.1539 | -0.6926 – 0.3849 | -0.5608 | 0.575 | 0.4681 | 0.0747 – 0.8615 | 2.3363 | **0.020** |  |  |  |  |  |  |  |  |
| Block * MB vs MW | -0.0476 | -0.6192 – 0.5240 | -0.1635 | 0.870 | -0.2247 | -0.6479 – 0.1985 | -1.0427 | 0.297 |  |  |  |  |  |  |  |  |
| Spont vs Delib MW |  |  |  |  |  |  |  |  | -1.1130 | -1.7421 – -0.4839 | -3.4740 | **0.001** | 0.7986 | 0.3371 – 1.2602 | 3.3978 | **0.001** |
| Block * Spont vs Delib MW |  |  |  |  |  |  |  |  | 0.2630 | -0.4147 – 0.9407 | 0.7621 | 0.446 | 0.0362 | -0.4671 – 0.5396 | 0.1414 | 0.888 |
| N | 31 Participant |  |  |  | 31 Participant |  |  |  | 31 Participant |  |  |  | 31 Participant |  |  |  |
| Marginal R<sup>2</sup> / Conditional R<sup>2</sup> | 0.002 / 0.110 |  |  |  | 0.014 / 0.300 |  |  |  | 0.018 / 0.132 |  |  |  | 0.021 / 0.313 |  |  |  |

MB vs. MW: Mind Blanking vs. Mind Wandering (higher scores refer to a tendency of mind wandering with accessible mental content). Spont vs Delib MW: Spontaneous vs. Deliberate mind wandering (higher scores reflect more deliberate mind wandering). LMMs were used with fixed slopes and random intercepts were used for participant due to convergence issues. Statistical parameters are based on blocks when participants reported mind wandering in contrast to task-focus comprising 646 observations in 31 participants.

### Slow periodic EEG activity is associated with both mind wandering and probabilistic learning

To examine the neural correlates of mind wandering and behavioral performance, we extracted the aperiodic component (the spectral exponent), and the periodic EEG components as measured during the blocks when participants actively engaged with the task. The spectral exponent, and the band-wise at each channel served as outcome variables in separate LMMs. First, we explored whether time as reflected by *Block* number was associated with changes in EEG activity, therefore, aperiodic and periodic EEG measures at each location were predicted by *Block* in separate LMMs.

As indicated in **Fig 5**, *Block* showed a considerable association with both aperiodic and periodic EEG activity. The slope of the EEG spectra showed a positive association with *Block* indicating a steepening of the EEG spectral slope as the task progressed. With regards to the periodic components, oscillatory activity in the slow and delta frequency range decreased, whereas activity in the theta, alpha and beta ranges increased as the task progressed. The most robust increase was evidenced in the alpha range.

**Figure 5.**
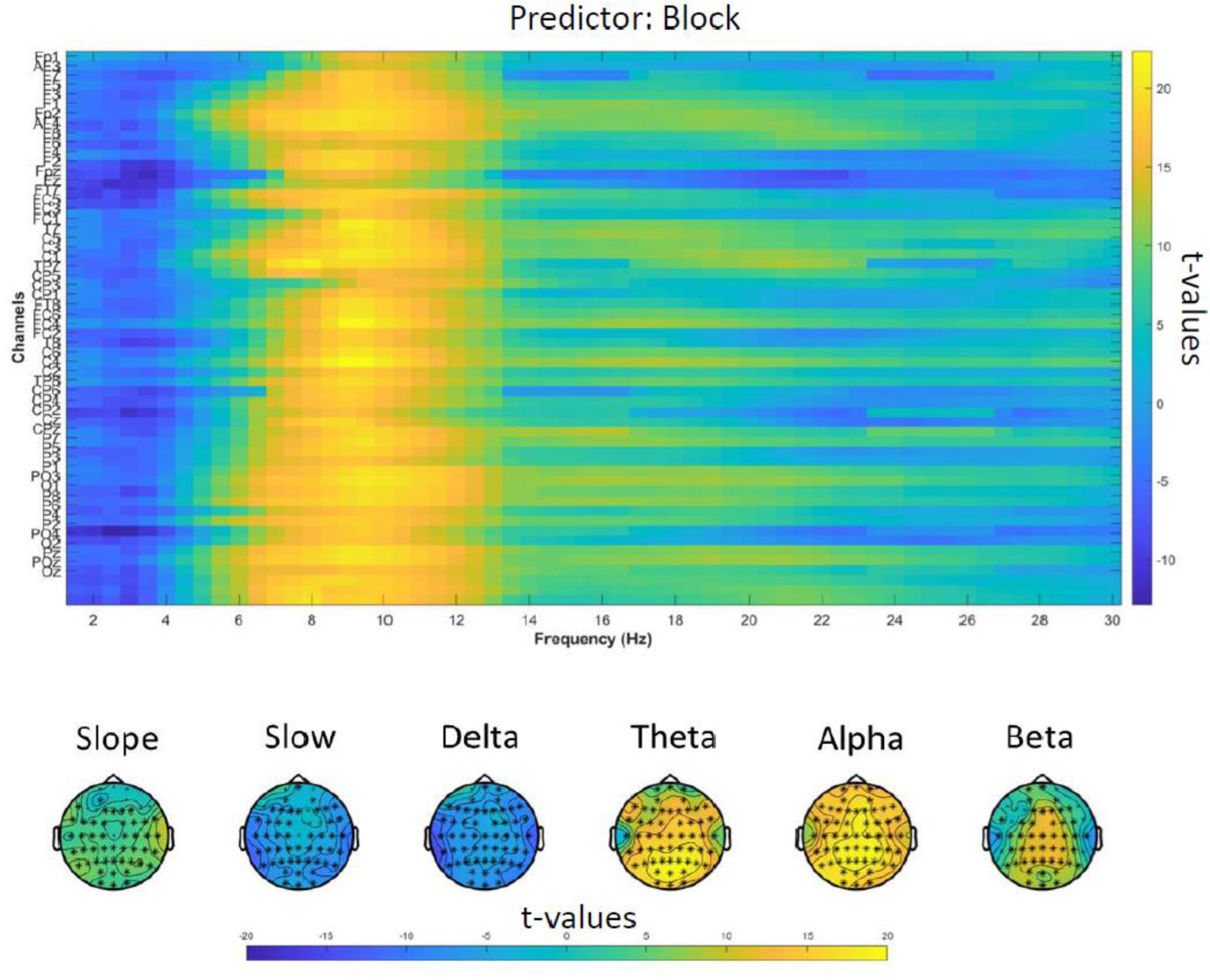
Aperiodic and periodic EEG activity predicted by Block number. LMMs predicting channel-wise EEG features by *Block* number indicate a robust change in EEG measures as the task progresses. The upper graph highlights the t-values of individual LMMs predicting oscillatory (periodic) EEG activity in each channel and frequency bin by *Block.* The lower headplots indicate the t-statistics of LMMs with aperiodic (Slope) and periodic activity (the later averaged over frequency bands) as outcome variables, and *Block* as the predictor. Asterisks on the headplots indicate statistically significant models after FDR correction. The findings indicate steepening spectral slope, decreasing slow and delta and increasing theta, alpha, beta oscillatory activity as the task progresses. The most pronounced increase occurred in the alpha (8-10 Hz) frequency range as indicated by the bin-wise analyses (upper graph).

Next, we examined the associations of *Mind Wandering* and *Probabilistic Learning* with EEG activity. Due to the robust effect of block number on EEG measures, we controlled for this confounding factor, and hence, performed LMMs predicting the spectral slope as well as the band-wise periodic EEG components by the fixed effects factors *Block* and *Mind Wandering* and by *Block* and *Probabilistic Learning* in separate models at each electrode location (including by-participant random effects). Given the distinct association between mind wandering and probabilistic learning as a function of time, we analyzed the first (1-15 blocks) and second half (16-30 blocks) of the task periods in separate models. Channel-wise analyses showed that in the first half of the task (1-15 blocks) periodic activity in the slow and delta frequency range was positively associated with both increased *Mind Wandering* and *Probabilistic Learning* peaking at centro-parietal sites, and to some extent at frontal and fronto-lateral locations. The association of slow and delta oscillatory activity with *Mind Wandering* and *Probabilistic Learning* showed considerable topographical overlaps (**Fig 6**). On the other hand, higher *Mind Wandering* was associated with a slight tendency (affecting only 4 electrode sites at frontopolar and right temporal locations) of a steepening spectral slope, and with a global increase in periodic activity in the beta range (comprising frontal, central and parietal sites), while increased *Probabilistic Learning* was associated with a tendency of a flattening slope (present in only one frontal electrode), and by decreased beta activity at right temporo-parietal locations.

**Figure 6.**
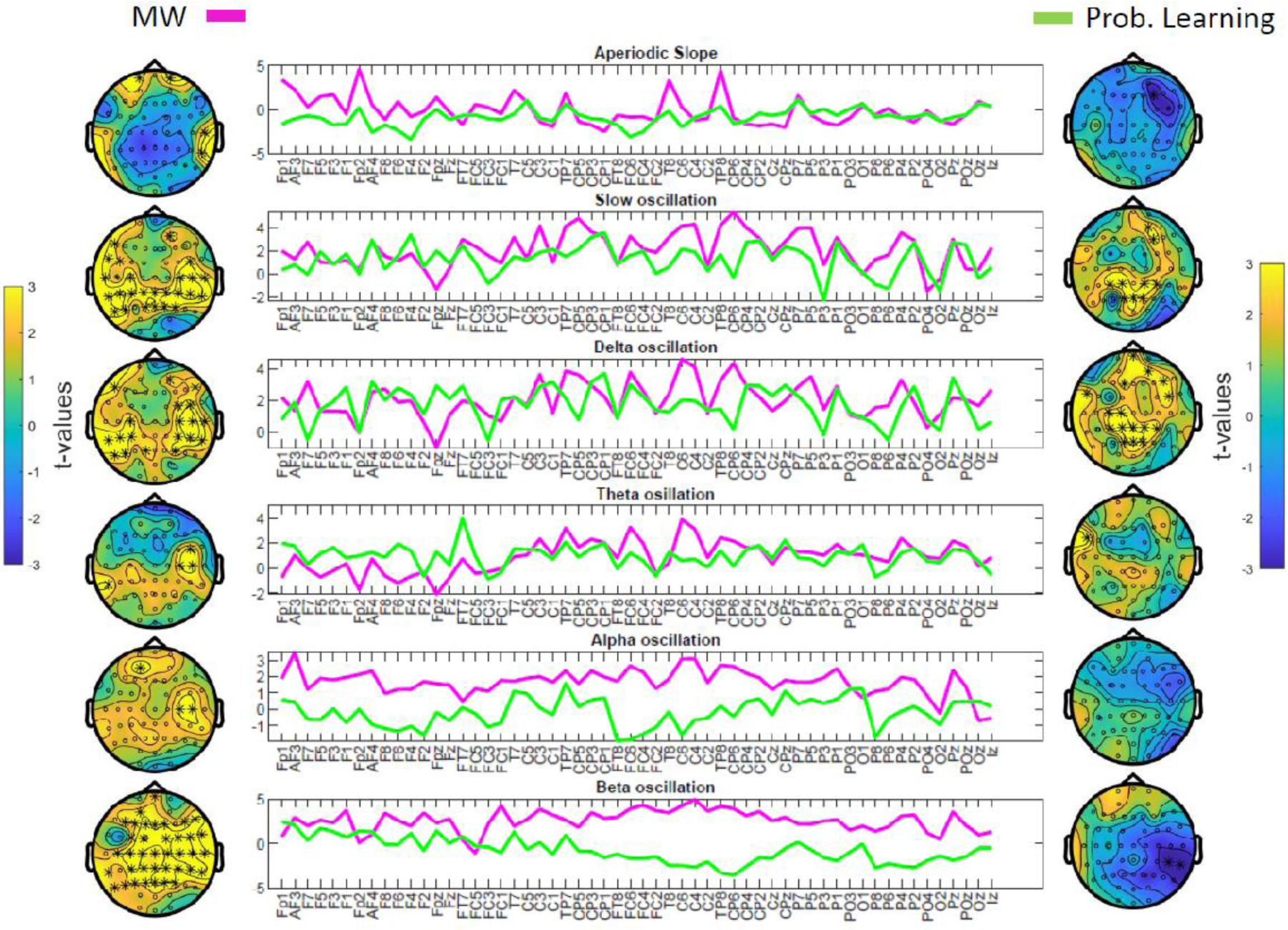
Topographical aspects of the associations of Mind Wandering and Probabilistic Learning with EEG activity in the first part (1-15 Block) of the ASRT task. LMMs predicting channel-wise EEG features by the independent variables *Block* and *Mind Wandering* as well as by *Block* and *Probabilistic Learning* were performed including the data of the first half of the task. *Mind Wandering* was associated with widespread oscillatory activity in the slow, delta, and beta frequency ranges, whereas *Probabilistic Learning* was linked to increased oscillatory activity in the slow and delta range. The t-statistics of the predictors *Mind Wandering* (left right, and magenta colored lineplots) and *Probabilistic Learning* (right side and green colored lineplots) are shown for each EEG feature and channel. Significant channels after FDR correction are highlighted by asterisks. *Mind Wandering* scores were inverted to facilitate understanding, that is, higher scores reflected higher mind wandering. Slow and delta oscillatory correlates of *Mind Wandering* and *Probabilistic Learning* showed considerable overlaps especially in centroparietal sites.

EEG correlates of mind wandering in the second half (16-30 blocks) of the task consisted of a right lateralized centroparietal increase in the spectral slope, and widespread increases in theta and alpha oscillatory activity over posterior regions, as well as an increase in beta oscillatory activity peaking over midline locations along the anterio-posterior axis. Probabilistic learning was not associated with changes in aperiodic or periodic EEG activity in the second half of the task (**Fig 7**).

**Figure 7.**
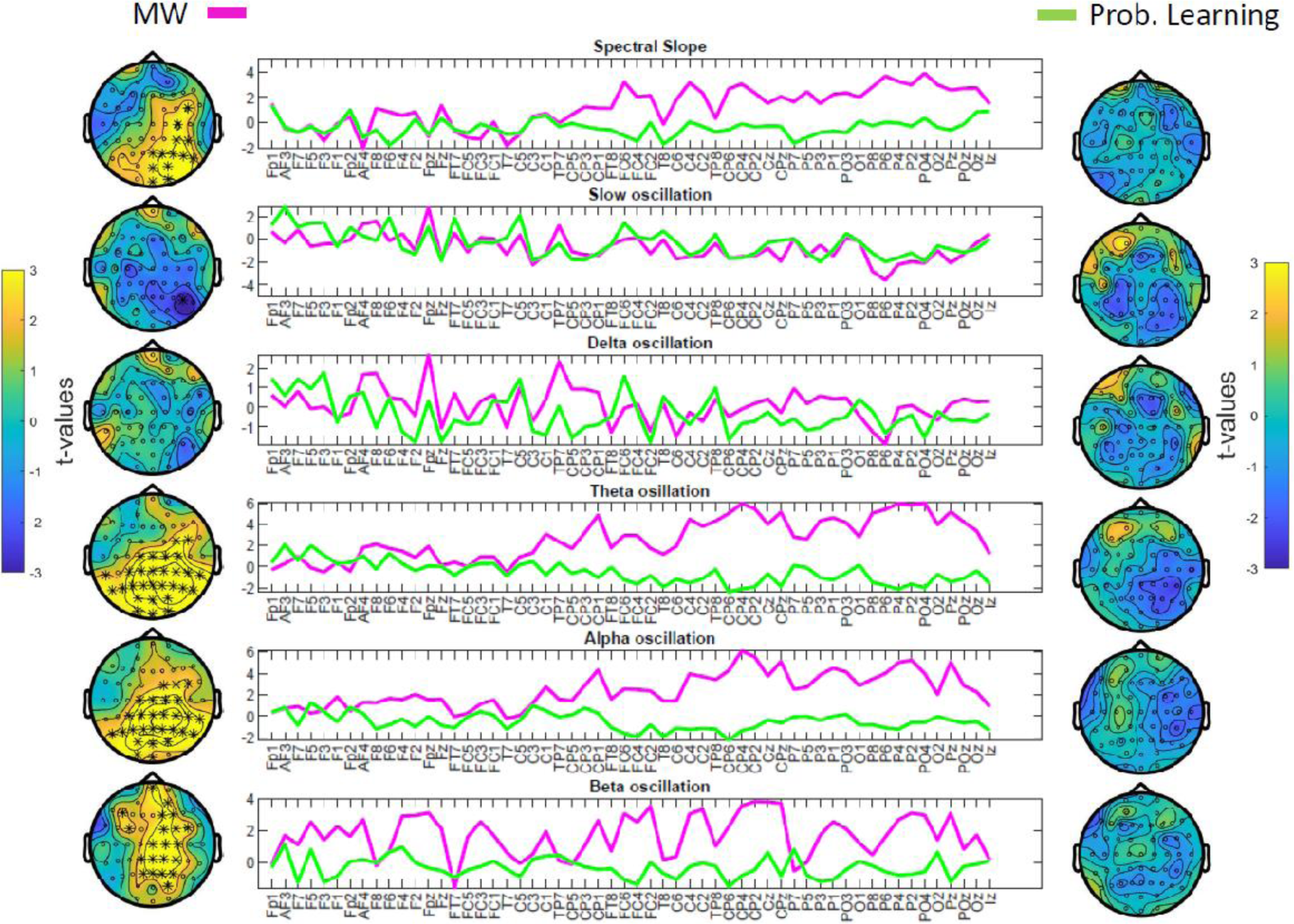
Topographical aspects of the associations of Mind Wandering and Probabilistic Learning with EEG activity in the second part (16-30 Block) of the ASRT task. LMMs predicting channel-wise EEG features predicted by *Block* and *Mind Wandering* as well as by *Block* and *Probabilistic Learning* were performed including the data of the second half of the task. *Mind Wandering* was associated with steeper spectral slope over right lateralized centroparietal sites and widespread increases in posterior theta and alpha, and midline beta oscillatory activity. *Probabilistic Learning* was not associated with changes in EEG features in the second half of the task. The t-statistics of the predictors *Mind Wandering* (left right, and magenta colored lineplots) and *Probabilistic Learning* (right side and green colored lineplots) are shown for each EEG feature and channel. Significant channels after FDR correction are highlighted by asterisks. *Mind wandering* scores were inverted to facilitate understanding, that is, higher scores reflected higher mind wandering.

## Discussion

A wide range of studies point to the harmful impact of mind wandering on everyday activities in natural settings (McVay et al., 2009a; Unsworth et al., 2012; Szpunar et al., 2013; Yanko and Spalek, 2014) and on cognitive processes measured in laboratory conditions (Mooneyham and Schooler, 2013). Such instances of task-unrelated thoughts often lead to erroneous responses, failure to detect targets and discard distracting items, or to a reduced ability to accurately encode, store or recall information (Miller, 2000; Braver, 2012; McVay and Kane, 2012b; Amer et al., 2016; Gratton et al., 2018; Andrillon et al., 2021; Blondé et al., 2022). Notably, the high prevalence of reports on the negative effects of mind wandering is due to studies focusing on attention-demanding cognitive operations, often defined in computational terms as model-based processes (Shohamy and Daw, 2014). Whereas model-based computations are cognitively more demanding, require executive control, and effortful, focused attention, model-free learning relies on acquiring habitual, and less attention-demanding stimulus-outcome dependencies (Daw et al., 2005; Otto et al., 2013; Pedraza et al., 2024). Here we showed that engaging in mind wandering might facilitate model-free processes, as exemplified by probabilistic learning. Our findings indicate that probabilistic learning—the ability to extract predictable patterns from a visual stream—was not only immune to the negative effects of mind wandering, but even appeared to benefit from such periods of inattention. Furthermore, improved probabilistic learning was linked to the nature of mind wandering: spontaneous, as opposed to deliberate mind wandering was associated with better probabilistic learning further supporting the link between transitory, uncontrolled lapses of attention and model-free computations. Finally, oscillatory activity in the low (slow and delta) frequency range indicative of covert sleep-like states was associated with both mind wandering and probabilistic learning in the first half of the task. Considering the key role of probabilistic learning in shaping behavior and underlying neural computations (Santolin and Saffran, 2018; Fiser and Lengyel, 2019), our findings add further insights into the benefits of task-unrelated thoughts in human cognition.

The benefit of mind wandering in probabilistic learning, may seem surprising in light of the large number of studies highlighting the costs of task-unrelated thoughts. Nevertheless, our finding does not stand alone without empirical antecedents. Emerging studies indicate that loosening attention and engaging in mind wandering or even entering into the transition between wakefulness and sleep onset might facilitate the processing hidden regularities in the perceptual information stream which are less accessible for model-based, goal-directed attention (Thompson-Schill et al., 2009; Amer et al., 2016; Decker et al., 2023; Vékony et al., 2023). Enhanced probabilistic learning in the ASRT task was previously shown to be associated with periods of mind wandering in an online study (Vékony et al, 2023). The present results, replicating, extending and validating this finding in a laboratory environment point to the robustness of this link. Of note, the implementation of the ASRT task in the current study was different from that of Vékony et al. (2023) in terms of stimulus type/spatial location, as well as pacing, suggesting that the beneficial effect of mind wandering on probabilistic learning generalizes across different contexts and task versions.

Better probabilistic learning under periods of mind wandering is in line with the assumptions of the competition framework suggesting an antagonistic relationship between model-based and model-free mental processes, or more specifically, between cognitive control and probabilistic learning (Poldrack and Packard, 2003; Janacsek et al., 2012; Pedraza et al., 2024). The competition framework postulates that probabilistic learning is enhanced under conditions of reduced executive control, an assumption that has received ample empirical support in recent years (Nemeth et al., 2013b; Virag et al., 2015; Ambrus et al., 2020; Pedraza et al., 2024). Since task-unrelated thoughts are consistently linked to reduced executive control (Smallwood et al., 2008; McVay et al., 2009b; McVay and Kane, 2010, 2012a; Kam and Handy, 2014; Kawagoe, 2022), we may speculate that improved probabilistic learning under periods of mind wandering might reflect an automatic shift to model-free from model-based forms of learning. By broadening the scope of attention, mind wandering might facilitate effortless and unconstrained exploration of the environment and promote favorable behaviors in situations where task rules and goals are somewhat opaque. The uncontrolled occurrence of task-unrelated thoughts might reflect such a shift in cognitive processes, and hence, facilitate the extraction of hidden (probabilistic) dependencies between stimuli.

Notably, mind wandering was linked to better probabilistic learning when it was experienced as spontaneous. Spontaneous task-unrelated thoughts seem to be different from deliberate mind wandering: the latter is characterized by the intention to engage in (usually) pleasant thoughts, feelings, and fantasies in a more constrained, goal-directed manner, while the former reflects the unintentional drifting of attention from the current task (Seli et al., 2015). Although the mechanisms differentiating spontaneous from deliberate mind wandering are not fully understood, spontaneous mind wandering is more consistently associated with executive failures and inattention, whereas engaging in deliberate mind wandering appears to be linked to reduced motivation to maintain task focus (Bozhilova et al., 2018; Robison and Unsworth, 2018; Robison et al., 2020). Interestingly, when compared to mind wandering with meta-awareness, episodes of task disengagement that remained unnoticed were associated with increased activity in several regions of the frontoparietal control network (Christoff et al., 2009), possibly providing an explanation for the stronger interference of spontaneously evolving mind wandering with neural mechanisms of task-related cognitive control.

Accordingly, our findings revealed distinct associations of the type of mind wandering with task performance. Whereas spontaneous mind wandering was linked to better probabilistic learning, deliberate mind wandering was associated with enhanced visuomotor accuracy. We may speculate that deliberate mind wandering was less detrimental to task performance as the task did not demand further attentional resources.

Mind wandering associated with inattention and deficits in executive control is thought to be triggered (at least partly) by transient, sleep-like activity expressed in a region-specific, localized manner (Andrillon et al., 2019; Jubera-Garcia et al., 2021). In fact, region-specific increases in slow waves, and increased slow frequency activity during attention demanding tasks were associated with both mind wandering and impaired behavioral performance (Andrillon et al., 2021; Wienke et al., 2021). However, these so-called offline states might not only have negative consequences on cognition and behavior. Offline states, accompanied by sleep-like cortical activity and experienced as mind wandering might facilitate the rapid consolidation of previously acquired information, similar to sleep-related memory consolidation. A large body of evidence has been accumulated during the last three decades regarding the beneficial influence of post-learning sleep on information processing (Diekelmann and Born, 2010; Rasch and Born, 2013). Sleep seems to be an ideal state for memory consolidation because of i) the attenuation of environmental stimuli and online encoding processes, ii) network activity at different neural levels (hippocampal, thalamocortical, cortico-cortical) facilitating the reactivation and reconsolidation of memories (Walker and Stickgold, 2010; Rasch and Born, 2013; Stickgold, 2013), and iii) optimization of signal to noise ratio by the readjustment of synaptic weights (Tononi and Cirelli, 2006). Although the idea that memories are specifically strengthened during sleep is intellectually appealing, recent studies question the exclusive role of sleep in consolidating previously encoded material, and suggest that periods of waking rest are equally beneficial for memory consolidation (Pan and Rickard, 2015; Wamsley, 2019, 2022; Németh et al., 2024; Ogawa et al., 2024).

Our findings, both at the behavioral and neural levels, suggest that the enhanced probabilistic learning observed during mind wandering may be driven by memory consolidation associated with local sleep in the waking brain (Wamsley and Summer, 2020; Wamsley, 2022). This assumption extends classical theories of memory consolidation, such as sleep-dependent and time-dependent memory consolidation, proposing a new type of consolidation termed as local-sleep-dependent consolidation (Vékony et al., 2023). Local-sleep-dependent consolidation in tasks requiring model-free learning may explain the inconsistent findings regarding sleep-dependent memory consolidation in diverse probabilistic and procedural learning tasks (Pan and Rickard, 2015; Németh et al., 2024). Our assumption aligns with the opportunistic memory consolidation theory, which suggests that consolidation can occur while awake, asleep (Mednick et al., 2011), or - as we suggest-under local-sleep states. Furthermore, local-sleep-dependent consolidation may not only stabilize past memories but also support the formation of predictive representations. This way, such transient offline periods may facilitate the extraction of regularities and underlying probabilities, which are essential to predictive processes. However, further studies employing magnetoencephalography or electroencephalography are needed to test this idea, providing direct evidence of the relationship between mind wandering, local sleep, as well as the formation and updating of predictive representations.

In sum, our findings suggest that the strengthening of memories acquired through model-free learning, such as implicit probabilistic learning, might not even require periods of waking rest, but could undergo rapid consolidation during task acquisition, especially when participants enter into a state of mind wandering (i.e. a transient offline state). The positive association between probabilistic learning and mind wandering during task acquisition may also explain why probabilistic learning occurs mainly through practice (Quentin et al., 2021), especially if post-practice rest periods are limited (Szücs-Bencze et al., 2023), and why probabilistic learning does not seem to benefit further from post-learning sleep or long periods of rest (Nemeth et al., 2010; Simor et al., 2019). Notably, in our study, the association between mind wandering and probabilistic learning showed an interaction with the time spent engaging with the task. The positive link between mind wandering and probabilistic learning was evidenced in the beginning, whereas it vanished at the second part of the task. The advantage of mind wandering over task focus, when participants start to practice the task, is in line with the assumption that mind wandering during task-acquisition may facilitate the extraction and rapid consolidation of probabilistic information hidden in the visual stream.

In line with the above, EEG correlates of mind wandering and probabilistic learning showed considerable topographical overlaps in the first half of the task. In particular, increases in slow and delta oscillatory activity, most prominently in centro-parietal locations (pointing to the involvement of sensorimotor regions (Melnik et al., 2017)) were linked to enhanced mind wandering and probabilistic learning. Since low frequency activity during sleep and waking rest seem to have key roles in offline memory consolidation, our observation points to the potential influence of covert sleep states on mind wandering and probabilistic learning. These findings are interesting because in the ASRT task, the raw probabilities and statistics are extracted from the visual information stream precisely in the first half of the task (Janacsek and Nemeth, 2012). Hence, slow and delta oscillatory activity, along with mind wandering, may support the discovery of probabilistic patterns within the information stream. Nevertheless, whether centro-parietal low frequency oscillations reflect covert and local sleep-like states over sensorimotor regions, and play a role in facilitating information processing remains to be explored. Future studies should systematically manipulate oscillatory activity and/or sleep pressure during task performance to infer the mechanistic role of oscillatory activity in mind wandering and information processing.

The EEG correlates of mind wandering, however, were not limited to low frequency oscillatory activity. In the first half of the task a widespread increase in beta oscillatory activity was observed. In addition, in the second half of the task mind wandering was associated with the aperiodic component, more specifically with steeper spectral slopes in posterior regions, as well as with increased centro-parietal theta and alpha, and an increase in the beta band in midline locations along the antero-posterior axis. Enhanced spectral power in the theta and alpha ranges were observed in previous studies as correlates of mind wandering (Kam et al., 2022). Interestingly, the EEG correlates of mind wandering in the second half of the task resemble the pattern observed in relation to the influence of time spent practicing the task (i.e. the effect of block number). A steeper spectral slope reflecting enhanced inhibition over excitation as the background activity, and increased periodic activity in theta-alpha range might reflect perceptual decoupling and a drift in attention towards self-generated thoughts. Shifting attention to the internal world from the external environment might be more prevalent when visuomotor control becomes more and more automatic as the task progresses. Oscillatory activity in the beta range was positively associated with mind wandering in both the first and the second half of the task. Interestingly, beta power was not consistently related to mind wandering in previous studies (exhibiting increases, decreases or null effects) (Rodriguez-Larios et al., 2021; Kam et al., 2022; Musat et al., 2024). We should note however, that it may seem difficult to compare the EEG correlates of mind wandering in tasks which differ in cognitive load, cognitive processes or focus of attention. That is, the phenomenological aspects, the neural correlates, as well as the functions of mind wandering might considerably differ in different contexts. Moreover, previous studies examining the EEG spectra during mind wandering mostly focused on band-wise spectral power overlooking and intermingling the aperiodic and periodic components of the EEG spectrum (Donoghue et al., 2020). Whereas the aperiodic component reflects background activity and neural excitability (i.e. excitation/inhibition balance, Gao et al., 2017), periodic components may capture moment-to-moment changes in neural firing in response to external stimuli or internally driven cognitive processes (Csercsa et al., 2010; Halgren et al., 2019; Ujma et al., 2022).

Despite its potential benefits, mind wandering was also associated with cognitive costs: loosening task focus was linked to poorer visuomotor performance, that is, participants in general (regardless of the probabilistic nature of the trials) committed more errors in blocks when they reported to mind wander. Such findings are in line with previous reports accentuating the negative aspects of mind wandering (Mooneyham and Schooler, 2013; Smallwood and Schooler, 2015), and suggesting that task-unrelated thoughts disrupt task performance (Andrillon et al., 2019, 2021). Whereas the costs of mind wandering in attention-demanding tasks are widely established, future studies combining attentional or executive control with probabilistic learning in a single task could examine whether the costs of mind wandering in attention and control also convey its benefits in other domains, such as probabilistic learning.

Our study is not without limitations. Since we assessed mind wandering only at the end of each block, our database contained a relatively limited number of self-reports. While increasing the number of thought-probes throughout the task would have been advantageous, frequent sampling during practice could heighten metacognition and interrupt the stream of perceptual stimuli, potentially interfering with task performance (He and Li, 2023). Moreover, although we explored the types of task-unrelated thoughts to some extent, the mental experience of mind wandering is evidently more complex and presumably way more heterogeneous. Future studies should assess the phenomenological aspects of mind wandering in more detail in order to disentangle the potential benefits and costs of task-unrelated thoughts on task performance.

Despite these limitations, our study provides empirical support for the potential benefits of mind wandering in information processing, and indicates that although visuomotor accuracy is reduced, probabilistic learning is enhanced during periods of attenuated task focus. Our study suggests that mind wandering and transient, local sleep-like states might facilitate the rapid extraction and consolidation of hidden regularities from the visual information stream. This phenomenon may provide the basis for the acquisition of new skills and optimization of predictive processes.

## Acknowledgements

This work was supported by the Chaire de Professeur Junior Program by INSERM and French National Grant Agency (ANR-22-CPJ1-0042-01); the National Brain Research Program project NAP2022-I-2/2022 (DN); Hungarian National Research, Development and Innovation Office Grant NKFI FK 142945 (PS); Janos Bolyai scholarship of the Hungarian Academy of Sciences (PS).

## Supplementary Materials

### S1. Instructions for thought probes

*“Throughout the task, try to stay focused on the task. However, you may find that your mind may wander, which is a perfectly natural phenomenon. We will interrupt the task a few times and ask three questions about it each time, whether you were concentrating on the task at hand or on something else. To answer these questions, use the mouse!*

*First, we ask you to what extent you focused on the task before the question appeared. Here, the answer options are on a 4-point scale from “1: Not at all” to “4: Completely”. A response of “1: Not at all” means that your thoughts are completely diverted (e.g. to your friends, weekend plans, etc.). A “4: Completely” answer means that you were only thinking about the task at hand (where the dog’s head appears, and which key to press as soon as possible)*.

*The second question asks whether, if you were not focused on the task, you thought about specifically about something, or if you thought about nothing at all. Here, the answer options are on a 4-point scale ranging from “1: I was thinking about nothing” to “4: I was thinking about something in particular”. A response of “1: I was thinking about nothing” means that your mind wandered, but you weren’t thinking about anything it’s as if your mind was completely blank. The answer “4: I was thinking about something in particular” means that you were thinking about something (e.g. a book, recent events, the task is too easy, you are hungry, it is uncomfortable to sit to do the task, etc.)*.

*The third question refers to whether your focus of attention (either on the task or elsewhere) was intentional or spontaneous. Here, the response options are on a 4-point scale from ‘1: I was completely spontaneous” to “4: I was completely deliberate” A response of “1 - I was completely spontaneous “ means that your attention did not require any effort. If you were focusing on the task at hand, it happened automatically, it was not difficult to focus. If you were distracted, this happened even though you wanted to focus on the task at hand. The answer “4: I was completely spontaneous “ means that you consciously directed your attention somewhere. If you were focusing on the task at hand, it was conscious, you were deliberately focusing on the task. If your mind was wandering, it was because you made a conscious decision to focus on something else. (e.g., the weekend). This can happen when you are bored during the task. It is important to stress that there is no right or wrong answer to these questions, and that your answers will not have any consequences. Please answer the questions completely honestly!”*

### S2. Quiz questions

The following quiz questions were presented to the participants following the instructions (see above “*Instructions for thought probes*”).

1. When doing the task, I thought of a film I saw yesterday:
2. I was completely focused on the task
3. Not focused on the task at all
4. If, while doing the task, I was thinking about my dinner tonight, I was:
5. Completely focused on the task
6. Not focused on the task at all
7. If, while doing the task, I was focused on where the dog’s head would appear, I was:
8. I focused completely on the task
9. I did not focus on the task at all
10. If I was focused on the noise in the room while doing the task, I was:
11. Focused completely on the task
12. Not focused on the task at all
13. If I was concentrating on which key to press while doing the task, then:
14. I focused completely on the task
15. I did not focus on the task at all
16. If, while doing the task, I was thinking about how easy it was or how difficult the task is, then:
17. I focused completely on the task
18. I did not focus on the task at all
19. If, while doing the task, I did not focus on the task but I was thinking about the task, then:
20. I was thinking specifically about something
21. I was not thinking about anything
22. If, while doing the task, I tried to focus on the task, but my thoughts wandered, then:
23. It was intentional
24. It was completely spontaneous

**S3.** Mind wandering and ASRT task performance. LMM random intercept models with RT based indices of *Probabilistic Learning* and *Visuomotor RT* as outcome variables, *Block* and *Mind Wandering* versus *Task Focus*, as well as their interaction as fixed effects predictors.

| *Predictors* | **Probabilistic Learning RT** |  |  |  | **Visuomotor RT** |  |  |  |
| --- | --- | --- | --- | --- | --- | --- | --- | --- |
|  | *Estimates* | *95 % CI* | *t-value* | *p* | *Estimates* | *95 % CI* | *t-value* | *p* |
| (Intercept) | 4.3291 | 2.3238 – 6.3344 | 4.2336 | <0.001 | 357.2747 | 347.4049 – 367.1445 | 70.9875 | <0.001 |
| Block | 0.0931 | 0.0067 – 0.1794 | 2.1127 | 0.035 | -0.0957 | -0.1750 – -0.0164 | -2.3680 | 0.018 |
| MW vs Task Focus | 0.6983 | -1.4789 – 2.8754 | 0.6289 | 0.529 | 0.0228 | -1.9872 – 2.0327 | 0.0222 | 0.982 |
| Block * MW vs Task Focus | 0.0290 | -0.0906 – 0.1486 | 0.4754 | 0.635 | 0.0066 | -0.1041 – 0.1172 | 0.1164 | 0.907 |
| ICC | 0.05 |  |  |  | 0.78 |  |  |  |
| N | 37 Participant |  |  |  | 37 Participant |  |  |  |
| Marginal R<sup>2</sup> / Conditional R<sup>2</sup> | 0.003 / 0.052 |  |  |  | 0.001 / 0.779 |  |  |  |

Block spans between 1 and 30. In the case of MW vs. Task Focus lower scores indicate relatively increased mind wandering as centered to the participant’s own mean.

**S4.**
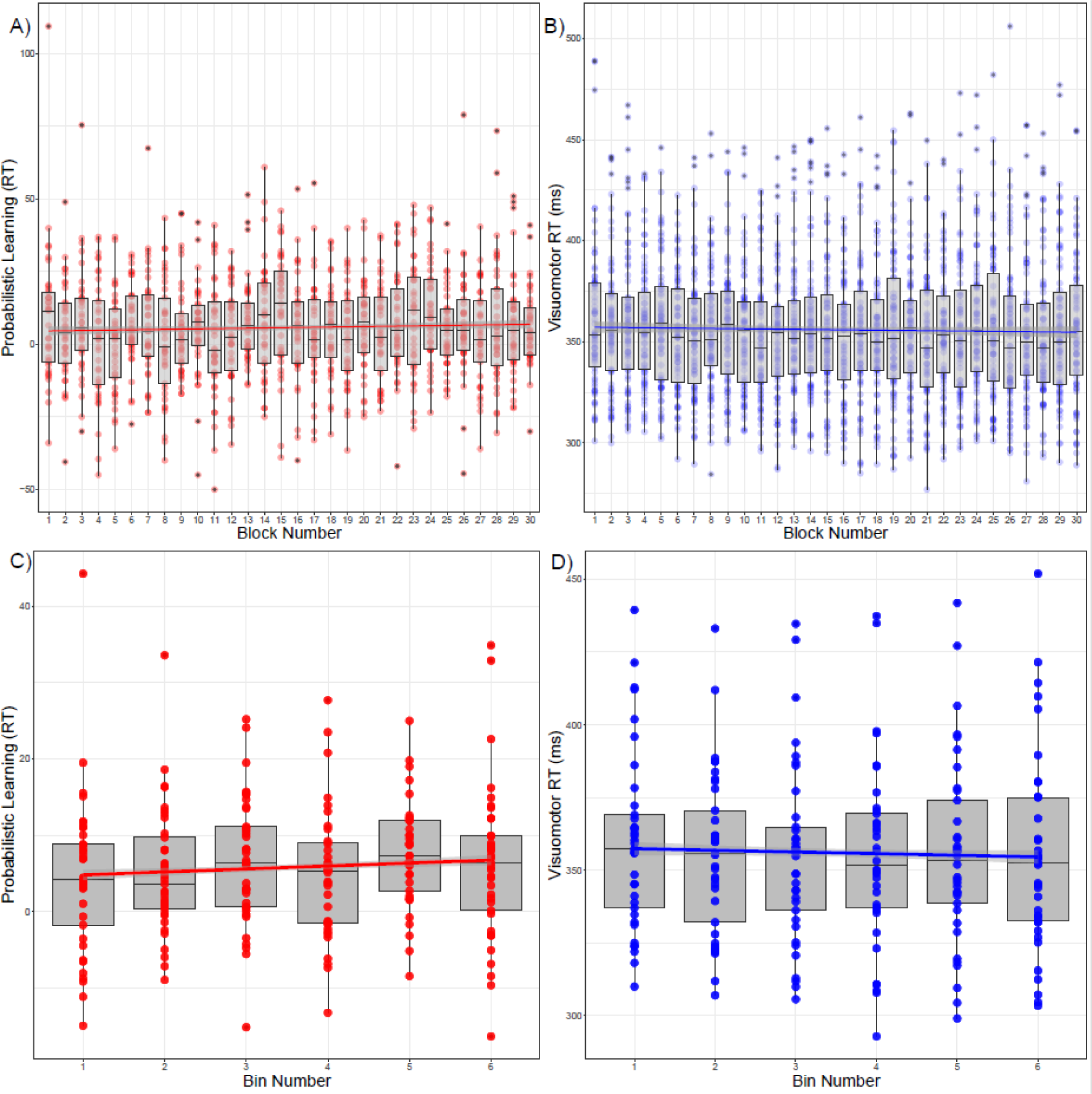
RT based Probabilistic Learning and Visuomotor Performance over successive blocks and bins. *Probabilistic Learning* is quantified by the difference in reaction times (medians for correct trials) between low-probability and high-probability trials averaged within each block (A, C). *Probabilistic Learning* gradually increases throughout the task due to relatively faster responses to high-versus low-probability trials. *Visuomotor RT* (response times regardless of trial types) (B, D) also changes throughout the task due to gradually shorter reaction times regardless of trial types. The upper graphs depict performance across each block. In the lower graphs (to facilitate visualization) the same performance is averaged over successive steps of five consecutive blocks (Blocks: 1-5, 6-10…26-30) marked as bins.

## References

Aasen SR, Drevland RN, Csifcsák G, Mittner M (2024) Increasing Mind Wandering With Accelerated Intermittent Theta Burst Stimulation Over the Left Dorsolateral Prefrontal Cortex. Available at: https://osf.io/fkx3w/download [Accessed July 18, 2024].

Alexandersen A, Csifcsák G, Groot J, Mittner M (2022) The effect of transcranial direct current stimulation on the interplay between executive control, behavioral variability and mind wandering: A registered report. Neuroimage: Reports 2:100109.

Allaire J (2012) RStudio: integrated development environment for R. Boston, MA 770:165–171.

Ambrus GG, Vékony T, Janacsek K, Trimborn AB, Kovács G, Nemeth D (2020) When less is more: Enhanced statistical learning of non-adjacent dependencies after disruption of bilateral DLPFC. Journal of Memory and Language 114:104144.

Amer T, Campbell KL, Hasher L (2016) Cognitive Control As a Double-Edged Sword. Trends Cogn Sci 20:905–915.

Andrillon T, Burns A, Mackay T, Windt J, Tsuchiya N (2021) Predicting lapses of attention with sleep-like slow waves. Nature Communications 12:1–12.

Andrillon T, Windt J, Silk T, Drummond SPA, Bellgrove MA, Tsuchiya N (2019) Does the Mind Wander When the Brain Takes a Break? Local Sleep in Wakefulness, Attentional Lapses and Mind-Wandering. Front Neurosci 13:949.

Aslin RN (2017) Statistical learning: a powerful mechanism that operates by mere exposure. WIREs Cognitive Science 8:e1373.

Benjamini Y, Hochberg Y (1995) Controlling the False Discovery Rate: A Practical and Powerful Approach to Multiple Testing. Journal of the Royal Statistical Society Series B: Statistical Methodology 57:289–300.

Blondé P, Girardeau J-C, Sperduti M, Piolino P (2022) A wandering mind is a forgetful mind: A systematic review on the influence of mind wandering on episodic memory encoding. Neuroscience & Biobehavioral Reviews 132:774–792.

Bódizs R, Szalárdy O, Horváth C, Ujma PP, Gombos F, Simor P, Pótári A, Zeising M, Steiger A, Dresler M (2021) A set of composite, non-redundant EEG measures of NREM sleep based on the power law scaling of the Fourier spectrum. Scientific reports 11:1–18.

Bonifacci P, Viroli C, Vassura C, Colombini E, Desideri L (2023) The relationship between mind wandering and reading comprehension: A meta-analysis. Psychon Bull Rev 30:40–59.

Bozhilova NS, Michelini G, Kuntsi J, Asherson P (2018) Mind wandering perspective on attention-deficit/hyperactivity disorder. Neuroscience & Biobehavioral Reviews 92:464–476.

Braver TS (2012) The variable nature of cognitive control: a dual mechanisms framework. Trends in Cognitive Sciences 16:106–113.

Brosowsky NP, Murray S, Schooler JW, Seli P (2021) Attention need not always apply: Mind wandering impedes explicit but not implicit sequence learning. Cognition 209:104530.

Christoff K, Gordon AM, Smallwood J, Smith R, Schooler JW (2009) Experience sampling during fMRI reveals default network and executive system contributions to mind wandering. Proceedings of the National Academy of Sciences 106:8719–8724.

Colombo MA, Napolitani M, Boly M, Gosseries O, Casarotto S, Rosanova M, Brichant J-F, Boveroux P, Rex S, Laureys S (2019) The spectral exponent of the resting EEG indexes the presence of consciousness during unresponsiveness induced by propofol, xenon, and ketamine. Neuroimage 189:631–644.

Csercsa R et al. (2010) Laminar analysis of slow wave activity in humans. Brain 133:2814–2829.

Daw ND, Niv Y, Dayan P (2005) Uncertainty-based competition between prefrontal and dorsolateral striatal systems for behavioral control. Nat Neurosci 8:1704–1711.

Decker A, Dubois M, Duncan K, Finn AS (2023) Pay attention and you might miss it: Greater learning during attentional lapses. Psychon Bull Rev 30:1041–1052.

Diekelmann S, Born J (2010) The memory function of sleep. Nature Reviews Neuroscience 11:114–126.

Donoghue T, Haller M, Peterson EJ, Varma P, Sebastian P, Gao R, Noto T, Lara AH, Wallis JD, Knight RT (2020) Parameterizing neural power spectra into periodic and aperiodic components. Nature neuroscience 23:1655–1665.

Drevland RN, Aasen SR, Csifcsák G, Mittner M (2024) Reducing mind wandering using continuous theta burst stimulation. Available at: https://osf.io/u5j7s/download [Accessed July 18, 2024].

Fiser J, Aslin RN (2001) Unsupervised Statistical Learning of Higher-Order Spatial Structures from Visual Scenes. Psychol Sci 12:499–504.

Fiser J, Lengyel G (2019) A common probabilistic framework for perceptual and statistical learning. Current Opinion in Neurobiology 58:218–228.

Gao R, Peterson EJ, Voytek B (2017) Inferring synaptic excitation/inhibition balance from field potentials. NeuroImage 158:70–78.

Gerster M, Waterstraat G, Litvak V, Lehnertz K, Schnitzler A, Florin E, Curio G, Nikulin V (2022) Separating Neural Oscillations from Aperiodic 1/f Activity: Challenges and Recommendations. Neuroinform 20:991–1012.

Gratton G, Cooper P, Fabiani M, Carter CS, Karayanidis F (2018) Dynamics of cognitive control: Theoretical bases, paradigms, and a view for the future. Psychophysiology 55:e13016.

Groot JM, Csifcsák G, Wientjes S, Forstmann BU, Mittner M (2022) Catching wandering minds with tapping fingers: neural and behavioral insights into task-unrelated cognition. Cerebral Cortex 32:4447–4463.

Halgren M, Ulbert I, Bastuji H, Fabó D, Erőss L, Rey M, Devinsky O, Doyle WK, Mak-McCully R, Halgren E, Wittner L, Chauvel P, Heit G, Eskandar E, Mandell A, Cash SS (2019) The generation and propagation of the human alpha rhythm. Proc Natl Acad Sci U S A 116:23772– 23782.

He H, Li H (2023) The Influence of Probe Frequency on Self-Reported Mind Wandering During Tasks With Different Cognitive Loads. Psychol Rep:00332941231214504.

Horváth K, Nemeth D, Janacsek K (2022) Inhibitory control hinders habit change. Scientific Reports 12:8338.

Howard JHJr, Howard DV (1997) Age differences in implicit learning of higher order dependencies in serial patterns. Psychology and aging 12:634.

Janacsek K, Fiser J, Nemeth D (2012) The best time to acquire new skills: age-related differences in implicit sequence learning across the human lifespan. Developmental Science 15:496–505.

Janacsek K, Nemeth D (2012) Predicting the future: from implicit learning to consolidation. International Journal of Psychophysiology 83:213–221.

Jubera-Garcia E, Gevers W, Van Opstal F (2021) Local build-up of sleep pressure could trigger mind wandering: Evidence from sleep, circadian and mind wandering research. Biochemical pharmacology 191:114478.

Kam JWY, Handy TC (2014) Differential recruitment of executive resources during mind wandering. Conscious Cogn 26:51–63.

Kam JWY, Rahnuma T, Park YE, Hart CM (2022) Electrophysiological markers of mind wandering: A systematic review. NeuroImage 258:119372.

Kawagoe T (2022) Executive failure hypothesis explains the trait-level association between motivation and mind wandering. Sci Rep 12:5839.

Kóbor A, Takács Á, Kardos Z, Janacsek K, Csépe V, Nemeth D (2018) ERPs differentiate the sensitivity to statistical probabilities and the learning of sequential structures during procedural learning. Biological psychology 135:180–193.

McVay JC, Kane MJ (2010) Does mind wandering reflect executive function or executive failure? Comment on Smallwood and Schooler (2006) and Watkins (2008). Psychological Bulletin 136:188–197.

McVay JC, Kane MJ (2012a) Drifting from slow to “d’oh!”: Working memory capacity and mind wandering predict extreme reaction times and executive control errors. Journal of Experimental Psychology: Learning, Memory, and Cognition 38:525.

McVay JC, Kane MJ (2012b) Why does working memory capacity predict variation in reading comprehension? On the influence of mind wandering and executive attention. Journal of experimental psychology: general 141:302.

McVay JC, Kane MJ, Kwapil TR (2009a) Tracking the train of thought from the laboratory into everyday life: an experience-sampling study of mind wandering across controlled and ecological contexts. Psychon Bull Rev 16:857–863.

McVay JC, Kane MJ, Kwapil TR (2009b) Tracking the train of thought from the laboratory into everyday life: An experience-sampling study of mind wandering across controlled and ecological contexts. Psychonomic Bulletin & Review 16:857–863.

Mednick SC, Cai DJ, Shuman T, Anagnostaras S, Wixted JT (2011) An opportunistic theory of cellular and systems consolidation. Trends in Neurosciences 34:504–514.

Melnik A, Hairston WD, Ferris DP, König P (2017) EEG correlates of sensorimotor processing: independent components involved in sensory and motor processing. Scientific Reports 7:4461.

Miller EK (2000) The prefontral cortex and cognitive control. Nat Rev Neurosci 1:59–65.

Mooneyham BW, Schooler JW (2013) The costs and benefits of mind-wandering: a review. Canadian Journal of Experimental Psychology/Revue canadienne de psychologie expérimentale 67:11.

Musat EM, Corcoran AW, Belloli L, Naccache L, Andrillon T (2024) Mind the blank: behavioral, experiential, and physiological signatures of absent-mindedness. bioRxiv:2024–02.

Németh D, Gerbier E, Born J, Rickard T, Diekelmann S, Fogel S, Genzel L, Prehn-Kristensen A, Payne J, Dresler M (2024) Optimizing the methodology of human sleep and memory research. Nature Reviews Psychology 3:123–137.

Nemeth D, Janacsek K, Fiser J (2013a) Age-dependent and coordinated shift in performance between implicit and explicit skill learning. Frontiers in Computational Neuroscience 7 Available at: https://www.frontiersin.org/article/10.3389/fncom.2013.00147 [Accessed May 6, 2022].

Nemeth D, Janacsek K, Londe Z, Ullman MT, Howard DV, Howard JH (2010) Sleep has no critical role in implicit motor sequence learning in young and old adults. Experimental brain research 201:351–358.

Nemeth D, Janacsek K, Polner B, Kovacs ZA (2013b) Boosting Human Learning by Hypnosis. Cerebral Cortex 23:801–805.

Nemeth D, Vékony T, Farkas B, Brezóczki B, Mittner M, Csifcsák G, Simor P (2023). Mind wandering enhances predictive processing.

Ogawa K, Yang Y, Yang H, Imai F, Imamizu H (2024) Human sensorimotor cortex reactivates recent visuomotor experience during awake rest. bioRxiv:2024–05.

Oldfield RC (1971) The assessment and analysis of handedness: The Edinburgh inventory. Neuropsychologia 9:97–113.

Otto AR, Raio CM, Chiang A, Phelps EA, Daw ND (2013) Working-memory capacity protects model-based learning from stress. Proceedings of the National Academy of Sciences 110:20941–20946.

Ouyang G, Hildebrandt A, Schmitz F, Herrmann CS (2020) Decomposing alpha and 1/f brain activities reveals their differential associations with cognitive processing speed. Neuroimage 205:116304.

Pan SC, Rickard TC (2015) Sleep and motor learning: is there room for consolidation? Psychological bulletin 141:812.

Pedraza F, Farkas BC, Vékony T, Haesebaert F, Phelipon R, Mihalecz I, Janacsek K, Anders R, Tillmann B, Plancher G (2024) Evidence for a competitive relationship between executive functions and statistical learning. npj Science of Learning 9:30.

Poldrack RA, Packard MG (2003) Competition among multiple memory systems: converging evidence from animal and human brain studies. Neuropsychologia 41:245–251.

Quentin R, Fanuel L, Kiss M, Vernet M, Vékony T, Janacsek K, Cohen LG, Nemeth D (2021) Statistical learning occurs during practice while high-order rule learning during rest period. npj Sci Learn 6:1–8.

Rasch B, Born J (2013) About sleep’s role in memory. Physiological reviews.

Robison MK, Miller AL, Unsworth N (2020) A multi-faceted approach to understanding individual differences in mind-wandering. Cognition 198:104078.

Robison MK, Unsworth N (2018) Cognitive and contextual correlates of spontaneous and deliberate mind-wandering. Journal of Experimental Psychology: Learning, Memory, and Cognition 44:85–98.

Rodriguez-Larios J, de Oca EABM, Alaerts K (2021) The EEG spectral properties of meditation and mind wandering differ between experienced meditators and novices. NeuroImage 245:118669.

Santolin C, Saffran JR (2018) Constraints on Statistical Learning Across Species. Trends Cogn Sci 22:52–63.

Schneider B, Szalárdy O, Ujma PP, Simor P, Gombos F, Kovács I, Dresler M, Bódizs R (2022) Scale-free and oscillatory spectral measures of sleep stages in humans. Front Neuroinform 16 Available at: https://www.frontiersin.org/articles/10.3389/fninf.2022.989262 [Accessed May 2, 2024].

Seli P, Carriere JS, Smilek D (2015) Not all mind wandering is created equal: Dissociating deliberate from spontaneous mind wandering. Psychological research 79:750–758.

Seli P, Konishi M, Risko EF, Smilek D (2018) The role of task difficulty in theoretical accounts of mind wandering. Consciousness and Cognition 65:255–262.

Shohamy D, Daw ND (2014) Habits and reinforcement learning. Available at: https://direct.mit.edu/books/edited-volume/chapter-pdf/2271330/c007100_9780262319362.pdf [Accessed April 24, 2024].

Simor P, Peigneux P, Bódizs R (2023) Sleep and dreaming in the light of reactive and predictive homeostasis. Neuroscience & Biobehavioral Reviews:105104.

Simor P, Zavecz Z, Horváth K, Ho N, Török C, Pesthy O, Gombos F, Janacsek K, Nemeth D (2019) Deconstructing procedural memory: Different learning trajectories and consolidation of sequence and statistical learning. Frontiers in Psychology 9:2708.

Smallwood J, Beach E, Schooler JW, Handy TC (2008) Going AWOL in the Brain: Mind Wandering Reduces Cortical Analysis of External Events. Journal of Cognitive Neuroscience 20:458–469.

Smallwood J, Davies JB, Heim D, Finnigan F, Sudberry M, O’Connor R, Obonsawin M (2004) Subjective experience and the attentional lapse: Task engagement and disengagement during sustained attention. Consciousness and Cognition 13:657–690.

Smallwood J, Schooler JW (2015) The Science of Mind Wandering: Empirically Navigating the Stream of Consciousness. Annual Review of Psychology:487–518.

Szpunar KK, Moulton ST, Schacter DL (2013) Mind wandering and education: From the classroom to online learning. Frontiers in Psychology 4.

Szücs-Bencze L, Fanuel L, Szabó N, Quentin R, Nemeth D, Vékony T (2023) Manipulating the rapid consolidation periods in a learning task affects general skills more than statistical learning and changes the dynamics of learning. Eneuro 10 Available at: https://www.eneuro.org/content/10/2/ENEURO.0228-22.2022.abstract [Accessed June 29, 2024].

Thompson-Schill SL, Ramscar M, Chrysikou EG (2009) Cognition without control: When a little frontal lobe goes a long way. Current directions in psychological science 18:259.

Tononi G, Cirelli C (2006) Sleep function and synaptic homeostasis. Sleep Medicine Reviews 10:49–62.

Ujma PP, Dresler M, Simor P, Fabó D, Ulbert I, Erőss L, Bódizs R (2022) The sleep EEG envelope is a novel, neuronal firing-based human biomarker. Sci Rep 12:18836.

Unsworth N, McMillan BD, Brewer GA, Spillers GJ (2012) Everyday attention failures: An individual differences investigation. Journal of Experimental Psychology: Learning, Memory, and Cognition 38:1765–1772.

Vékony T, Ambrus GG, Janacsek K, Nemeth D (2022) Cautious or causal? Key implicit sequence learning paradigms should not be overlooked when assessing the role of DLPFC (Commentary on Prutean et al.). Cortex 148:222–226.

Virag M, Janacsek K, Horvath A, Bujdoso Z, Fabo D, Nemeth D (2015) Competition between frontal lobe functions and implicit sequence learning: evidence from the long-term effects of alcohol. Exp Brain Res 233:2081–2089.

Wamsley EJ (2019) Memory consolidation during waking rest. Trends in cognitive sciences 23:171– 173.

Wamsley EJ (2022) Offline memory consolidation during waking rest. Nature Reviews Psychology 1:441–453.

Wamsley EJ, Summer T (2020) Spontaneous Entry into an “Offline” State during Wakefulness: A Mechanism of Memory Consolidation? Journal of Cognitive Neuroscience 32:1714–1734.

Wang LP, Maxwell SE (2015) On disaggregating between-person and within-person effects with longitudinal data using multilevel models. Psychol Methods 20:63–83.

Waschke L, Donoghue T, Fiedler L, Smith S, Garrett DD, Voytek B, Obleser J (2021) Modality-specific tracking of attention and sensory statistics in the human electrophysiological spectral exponent Chait M, Shinn-Cunningham BG, Postle BR, Simon JZ, eds. eLife 10:e70068.

Wienke C, Bartsch MV, Vogelgesang L, Reichert C, Hinrichs H, Heinze H-J, Dürschmid S (2021) Mind-wandering Is Accompanied by Both Local Sleep and Enhanced Processes of Spatial Attention Allocation. Cerebral Cortex Communications 2:tgab001.

Yanko MR, Spalek TM (2014) Driving With the Wandering Mind: The Effect That Mind-Wandering Has on Driving Performance. Hum Factors 56:260–269.

